# Stellate cell computational modelling predicts signal filtering in the molecular layer circuit of cerebellum

**DOI:** 10.1101/2020.08.25.266429

**Authors:** Martina Francesca Rizza, Francesca Locatelli, Stefano Masoli, Diana Sánchez Ponce, Alberto Muñoz, Francesca Prestori, Egidio D’Angelo

**Author notes:** **Correspondence:** Egidio D’Angelo.

## Abstract

The functional properties of cerebellar stellate cells and the way they regulate molecular layer activity are still unclear. We have measured stellate cells electroresponsiveness and their activation by parallel fiber bursts. Stellate cells showed intrinsic pacemaking, along with characteristic responses to depolarization and hyperpolarization, and showed a marked short-term facilitation during repetitive parallel fiber transmission. Spikes were emitted after a lag and only at high frequency, making stellate cells to operate as delay-high-pass filters. A detailed computational model summarizing these physiological properties allowed to explore different functional configurations of the parallel fiber – stellate cell – Purkinje cell circuit. Simulations showed that, following parallel fiber stimulation, Purkinje cells almost linearly increased their response with input frequency but such an increase was inhibited by stellate cells, which leveled the Purkinje cell gain curve to its 4 Hz value. When reciprocal inhibitory connections between stellate cells were activated, the control of stellate cells over Purkinje cell discharge was maintained only at very high frequencies. These simulations thus predict a new role for stellate cells, which could endow the molecular layer with low-pass and band-pass filtering properties regulating Purkinje cell gain and, along with this, also burst delay and the burst-pause responses pattern.

## Introduction

The stellate cells (SCs) are inhibitory interneurons of the cerebellum molecular layer (ML) that were first identified in histological preparations by Golgi (Golgi, 1874) and Cajal (Cajal, 1888, 1894). Electrophysiological recordings *in vivo* (Eccles, 1967) revealed their inhibitory nature. Since then, several observations have been reported on SC morphology and molecular properties but the way SCs control cerebellar functioning remained elusive.

SCs populate the outer two thirds of the cerebellum ML and provide inhibition to Purkinje cells (PCs) (Chan-Palay and Palay, 1972). Their dendrites are organized isotropically around the soma and lay on the parasagittal plane, where they are intercalated with the surrounding PC dendrites. The axon travels a short distance along the transverse plane, following the parallel fibers (PFs), and terminates with inhibitory synapses on PC dendrites as well as on other SCs generating feed-forward inhibition (Eccles, 1967; Midtgaard, 1992; Hausser and Clark, 1997; Jaeger and Bower, 1999; Mittmann et al., 2005; Barmack and Yakhnitsa, 2008; Rieubland et al., 2014). Several studies have suggested that, in combination with the direct excitatory pathway provided by PF - PC synapses, feed-forward inhibition may assure effective motor performance (Jaeger et al., 1997; Jaeger and Bower, 1999; Walter et al., 2006; Santamaria et al., 2007; Wulff et al., 2009; Bower, 2010) and learning (Jorntell et al., 2010; ten Brinke et al., 2015). Inhibition provided by ML interneurons proved also able to regulate the bandwidth of PC responses in spots and beams (Cohen and Yarom, 2000; Mapelli et al., 2010). Although the importance of SCs, there is little knowledge on their mechanisms of action (for a recent review see (Prestori et al., 2019)).

The electrophysiological properties of SCs have been identified only in part. SCs fire spontaneously in the 1–35 Hz range, both *in vitro* and *in vivo* (Llano and Gerschenfeld, 1993; Hausser and Clark, 1997; Carter and Regehr, 2002; Jorntell and Ekerot, 2002; Barmack and Yakhnitsa, 2008). The firing rate is dynamically regulated by several ionic channels, including T-type Ca^2+^ channels and A-type K^+^ channels (Molineux et al., 2005; Anderson et al., 2013; Alexander et al., 2019). Evidence has also been provided about the receptors and currents involved in synaptic transmission (Astori and Köhr, 2008). Given that a full characterization of all the relevant mechanisms is unavailable at the moment, a first hypothesis on stellate cell role could be generated through computational modeling, which can synthesize morpho-electrical information in a coherent rule-based framework (e.g. see the case of granule cell (GrC), Golgi cells and PCs, just to remain in the cerebellum). (D’Angelo et al., 2001; Solinas et al., 2007b, a; Diwakar et al., 2009; Masoli et al., 2015; Masoli and D’Angelo, 2017; Masoli et al., 2017; Masoli et al., 2020).

Initial attempts at modeling ML functions involved simplified representations of ML interneurons. Those models have shown the impact of increasing or decreasing SC transmission strength under the assumption of linear rate coding and in the absence of short-term plasticity (Santamaria et al., 2002; Lennon et al., 2014; Lennon et al., 2015). However, it is now clear from several studies that detailed membrane and synaptic dynamics are critical to understand the way a neural circuit operates (De Zeeuw et al., 2011; Arlt and Häusser, 2020). Therefore, we have recorded electroresponsive and synaptic dynamics of SC and used them to generate detailed computational models through well-defined workflows for model construction and validation (Druckmann et al., 2007; Masoli et al., 2017; Migliore et al., 2018; Masoli et al., 2020). The models, based on accurate morphologies and membrane mechanisms, faithfully reproduced the whole set of available experimental data. Importantly, a marked short-term facilitation at PF – SC synapses, as well as inhibitory transmission from other SCs, proved to be instrumental to modulate SC response curves to input bursts. The SC properties reverberated onto the PCs generating low-pass and band-pass filtering of PF signals. These observations imply that SCs transform the ML in a filter that limits the response bandwidth of on-beam PCs into the low-frequency range, leading to new hypotheses on how the cerebellar cortex processes incoming signals.

## Methods

### Electrophysiological recordings

All experiments were conducted in accordance with European guidelines for the care and use of laboratory animals (Council Directive 2010/63/EU), and approved by the ethical committee of Italian Ministry of Health (628/2017-PR). We performed whole-cell patch-clamp recordings (WCR) from SCs in acute cerebellar slices from P18–P25 male and female C57/BL6 mice. Briefly, mice were killed by decapitation after deep anesthesia with halothane (Sigma-Aldrich). The cerebellum was gently removed, and the vermis was isolated and fixed on the stage of a vibroslicer with cyanoacrylic glue. Acute 220-μm-thick slices were cut in the coronal plane in a cold (2–3 °C) oxygenated bicarbonate-buffered saline solution (Kreb’s solution) and maintained at room temperature for at least 1h, before being transferred to a recording chamber. The slices were continuously perfused at a rate of 1.5 ml/min with oxygenated Kreb’s solution and maintained at 32°C with a Peltier feedback device (TC-324B, Warner Instrument). The Kreb’s solution contained the following (in mM): NaCl 120, KCl 2, MgSO_4_ 1.2, NaHCO_3_ 26, KH_2_PO_4_ 1.2, CaCl_2_ 2, glucose 11 (pH 7.4 when equilibrated with 95%O_2_-5%CO_2_). SR 95531 (gabazine; 10 μM, Abcam) and strycnine (1 μM, Abcam) were added to the bath solution in order to block inhibitory synaptic inputs. Slices were visualized in an upright epifluorescence microscope (Axioskop 2 FS, Zeiss) equipped with a ×63, 0.9 NA water-immersion objective. Patch pipettes were fabricated from thick-walled borosilicate glass capillaries (Sutter Instruments) by means of a Sutter P-1000 horizontal puller (Sutter Instruments). Recordings were performed using a Multiclamp 700B [-3dB; cutoff frequency (fc), 10 kHz], sampled with Digidata 1550 interface, and analyzed off-line with pClamp10 software (Molecular Devices). SCs were recorded in the outer two-thirds of the molecular layer, where they are the only neuronal species present (Chan-Palay and Palay, 1972; Palay and Chan-Palay, 1974; Alcami and Marty, 2013).

PF stimulation was performed with a large-bore patch pipette filled with Kreb’s solution and placed across the molecular layer ~200 μm from the recording electrode. The stimulus intensity ranged from 10 to 50 V with duration of 0.2 ms. In order to analyze short-term dynamics during repetitive stimulation, trains of 10-20 pulses @ 50, 100 and 200 Hz were applied. All data are reported as mean ± SEM. Means were compared by a Student’s *t*-test or by one-way parametric analysis of variance (ANOVA). Where appropriate, data were further assessed by conducting the Tukey *post hoc* test. The analysis was two-sided, with level of significance α = 0.05.

#### Whole-cell recording properties

The stability of WCR can be influenced by modification of series resistance (R_s_). To ensure that R_s_ remained stable during recordings, passive electrode-cell parameters were monitored throughout the experiments. In each recording, once in the whole-cell configuration, the current transients elicited by 10 mV hyperpolarizing pulses from the holding potential of −70 mV in voltage-clamp mode showed a biexponential relaxation. Membrane capacitance (C_m_ = 6.7 ± 0.5 pF; n = 23) was measured from the capacitive charge (the area underlying current transients) and series resistance (R_s_ = 27.1 ± 3.1 MΩ; n = 23) was calculated as R_s_ = τ_VC_/C_m_. The input resistance (R_In_ = 1.3 ± 0.1 GΩ; n = 23) was computed from the steady-state current flowing after termination of the transient. The 3-dB cutoff frequency of the electrode-cell system, *f*_VC_ (1.2 ± 0.1; n = 23), was calculated as *f*_VC_ = (2π ∙ 2τ_VC_)^−1^. Series resistance was constantly monitored throughout the experiment. In analyzed recording periods, R_s_ was constant within 20%.

#### Stellate cell excitability

Pipette had a resistance of 7-10 MΩ resistance before seal formation with a filling solution containing the following (in mM): potassium gluconate 126, NaCl 4, HEPES 5, glucose 15, MgSO_4_ • 7H_2_O 1, BAPTA-free 0.1, BAPTA-Ca^2+^ 0.05, Mg^2+^-ATP 3, Na^+^-GTP 0.1, pH adjusted to 7.2 with KOH. Just after obtaining the cell-attached configuration, electrode capacitance was carefully cancelled to allow for electronic compensation of pipette charging during subsequent current-clamp recordings. After switching to current clamp, intrinsic excitability was investigated by setting the holding current at 0 pA and injecting 2 s steps of current (from −16 to 20 pA in a 4 pA increment). Action potential threshold (AP_thr_) was measured along the raising phase of membrane potential responses to step current injection. The AP_thr_ was identified at the flexus starting the regenerative process (a procedure that was improved by taking the second time derivative of membrane potential). Action potential duration at half amplitude (AP_HW_) was measured at the midpoint between the threshold and the peak. The amplitude of the action potential was estimated as the difference between the threshold and the maximum reached potential (AP_Ampl_). The amplitude of the action potential afterhyperpolarization (AP_AHP_) was estimated as the difference between threshold and the lowest potential after the peak. Action potential frequency (AP_Freq_) was measured by dividing the number of spikes by step duration and spikes times were used to calculated interspike intervals (ISI). Peri-stimulus time histograms (PSTHs) were constructed for the analysis of responses to 100 Hz-10 pulses stimulation. To optimize PSTH resolution, a 40 ms bin width was used.

#### Synaptic currents

Pipette had a resistance of 4-6 MΩ resistance before seal formation with a filling solution containing the following (in mM): 81 Cs_2_SO_4_, 4 NaCl, 2 MgSO_4_, 1 QX-314 (lidocaine *N*-ethyl bromide), 0.1 BAPTA-free, 0.05 BAPTA-Ca^2+^, 15 glucose, 3 Mg^2+^-ATP, 0.1 Na^+^-GTP and 15 HEPES, pH adjusted to 7.2 with CsOH. SCs were voltage-clamped at −70 mV. The EPSC_S_ were elicited at three types of high-frequency trains (20 pulses @ 50, 100 and 200 Hz). Each train stimulation sweep was separated from the preceding one by at least 30 s. EPSC amplitude value was calculated as the difference between peak and base. The facilitation during the repetitive stimulation was computed, for each Δt, by normalizing the amplitudes to the first EPSC.

### Tissue preparation for morphological reconstructions

For morphological reconstruction of molecular layer interneurons, adult C57/BL6 adult mice (8 weeks) were used. They received an overdose of sodium pentobarbital (0.09 mg/g, i.p.) and were perfused transcardially with phosphate-buffered saline (0.1 M PBS) followed by 4% paraformaldehyde in 0.1 M PB. Their brains were then removed and postfixed in 4% paraformaldehyde for 24 h. Vibratome parasagittal sections (200 μm) of the cerebellum were obtained. Sections were prelabeled with 4, 6-diamidino-2-phenylindole (DAPI; Sigma, St Louis, MO), and a continuous current was used to inject with Lucifer yellow (8 % in 0.1; Tris buffer, pH 7.4; LY) individual cells at different distances from the pial surface in the molecular layer of the cerebellum. Following the intracellular injections, sections were processed for immunofluorescence staining. They were incubated for 72 h at 4ºC in stock solution (2% bovine serum albumin, 1% Triton X-100, and 5% sucrose in PB) containing rabbit anti-LY (1:400 000; generated at the Cajal Institute). The sections were then rinsed in PB and incubated in biotinylated donkey anti-rabbit IgG (1:100; Amersham, Buckinghamshire, United Kingdom). Then, sections were then rinsed again and incubated Alexa fluor 488 streptavidin-conjugated (1:1000; Molecular Probes, Eugene, OR, United States of America). Finally, the sections were washed and mounted with ProLong Gold Antifade Reagent (Invitrogen Corporation, Carlsbad, CA, USA).

For cell reconstruction and quantitative analysis, sections were imaged with a confocal scanning laser attached to a fluorescence microscope Zeiss (LSM710). Consecutive stacks of images at high magnification (×63 glycerol; voxel size, 0.057 × 0.057 × 0.14 μm^3^ for mouse cells) were acquired to capture dendritic, and in some cases also axonal, arbors on the basis of Lucifer yellow immunostaining.

Data points of neuron morphology of each molecular layer interneuron were extracted in 3D using Neurolucida 360 (MicroBrightfield). Briefly, dendrites, axon and soma, in the skeleton definition were described through 3D points, delimiting the different segments that form the cell arbor. These points have an associated diameter that provides the information of the varying thickness of the dendritic or axonal processes at that particular point, and varies along the length of the processes. The soma was defined through a set of connected points tracing the contour of the soma in 2D. Morphological variables were extracted using Neurolucida software.

### Computational modeling

In this work, we have constructed and simulated multi-compartmental SC models, using Python-NEURON (Python 2.7.15; NEURON 7.7) (Hines and Carnevale, 2001; Hines et al., 2009). The models were based on four mouse SC detailed morphologies. The ionic channels were distributed over the somatic, dendritic and axonal compartments, according to immunohistochemical, electrophysiological and pharmacological data, and modeled following the HH formulation or Markov-chains for multi-state transitions, using mathematical methods reported previously (D’Angelo et al., 2001; Nieus et al., 2006; Solinas et al., 2007b, a; Anwar et al., 2012; Masoli et al., 2015).

The models were optimized with BluePyOpt (Van Geit et al., 2016), at the end of the optimization process the correct models are provided by the fine interaction of 14 ionic channels for a total of 36 maximum ionic conductances (G_i-max_) parameters assigned in the different morphology sections (Supplementary Fig. 1 and Table 1). G_i-max_ parameters have been found by an automatic optimization procedure to obtain the ensemble of electrophysiological behaviours. Each channel needed a precise value of G_i-max_ to interact, in an optimal way, with all the other values to achieve the exact balance between the sophisticate ionic channel dynamics.

#### Model morphology

Four mice stellate neurons were chosen from fluorescent images obtained with a confocal microscope and reconstructed with Neurolucida 360. The morphologies consisted of branched dendritic trees, a soma, and a branched axon with collaterals. The table (Supplementary Table 2) shows the number of morphological compartments, their diameter and length.

#### Passive properties and ionic channels

The passive properties of the SC model include R_axial_ which was set to 110 Ωcm for all the compartments, R_input_ = 0.75 ± 0.06 GΩ, C_m_ was set to 1.5 μF/cm^2^ for the dendrites while the rest of the compartments was set to 1μF/cm^2^. The reversal potential (E_rev_) of each ionic species was defined as follows: E_Na_ = 60 mV, E_k_ = −84 mV, E_Ca_ = 137.5 mV, E_h_ = −34 mV, E_leak_ = −48 mV. The leakage G_i-max_ was set to 3e-5 S/cm^2^ for all the compartments. The SC ionic channel models and distributions were taken from previous papers and updated according to the latest literature when needed (see Supplemental Material for details; (Masoli et al., 2015)). The model included Na^+^ channels (Nav1.1 and Nav1.6), K^+^ channels (Kv3.4, Kv4.3, Kv1.1, Kir2.3, Kv7, KCa1.1 and KCa2.2), Ca^2+^ channels (Cav2.1, Cav3.2 and Cav3.3), hyperpolarization-activated cyclic nucleotide–gated channels (HCN1) and a parvalbumin-based Ca^2+^ buffer.

#### Modeling excitatory and inhibitory neurotransmission

SCs receive synaptic activity from excitatory pathways, such as the GrC axons and the diffusion from climbing fibers and local inhibitory pathways. The inhibitory activity of SC modules the spike transmission to PC. Several studies have proved the presence of specific types of AMPA and NMDA receptor-mediated currents in cerebellar SC (Carter and Regehr, 2000; Nieus et al., 2006; Bidoret et al., 2015). AMPA and NMDA receptors, built with the Tsodyks formalism (Tsodyks et al., 1998), were used to simulate the excitatory synaptic activity from the PFs to SC dendrites.

The AMPA receptor received the following parameters (Nieus et al., 2006): release probability (*p*) = 0.15, recovery time constant (τ_R_) = 35.1 ms, facilitation time constant (τ_F_) = 10.8 ms, maximum ionic conductance (G_i-max_) = 2300 pS per synapse, ionic reversal potential (E_rev_) = 0 mV (Supplementary Fig. 2A and Table 3).

The NMDA kinetic scheme was adapted to simulate the presence of the NR2B subunit (Bidoret et al., 2015). NMDA NR2B receptor was modeled following the kinetics of the work of (Santucci and Raghavachari, 2008) and the fitted parameters were: *p* = 0.15, τ_R_ = 8 ms, τ_F_ = 5 ms, G_i-max_ = 10000 pS per synapse, E_rev_ = −3.7 mV (Supplementary Fig. 2B and Table 3).

The GABA_A_ receptor model maintains the kinetic scheme described in the work of (Nieus et al., 2014) and modified by maintaining the α1 subunit but deleting the α6 subunit (absent in the MLI) as follows: *p* = 0.42, τ_R_ = 38.7 ms, τ_F_ = 0 ms, G_i-max_ = 1600 pS per synapse, E_rev_ = −65 mV (Supplementary Fig. 2C and Table 3).

AMPA and NMDA receptors were placed on 3 distal dendrite compartments. GABA_A_ receptors were activated randomly on the dendrite compartments.

Model EPSCs and IPSCs were adapted to reproduce unitary synaptic current from SCs (Llano and Gerschenfeld, 1993; Nieus et al., 2006; Nieus et al., 2014) (Supplementary Fig. 2). The glutamatergic AMPA and NMDA receptors synaptic G_i-max_ parameter was balanced to reproduce a single PF EPSC, while *p* parameter was balanced to reproduce short-term facilitation in SC EPSCs during a stimulus train.

#### Optimization procedure

In this work, we used an innovative technique that allows rapid and automatic parameter optimization, based on multi-objective evolutionary algorithms, called “Indicator-Based Evolutionary Algorithm” (IBEA) (Zitzler and Kunzli, 2004), in the BluePyOpt Framework (Van Geit et al., 2016), to estimate the maximum ionic conductances parameters (G_i-_ max) (the free parameters of the model) of all ionic channels distributed along the morphology sections obtaining models able to reproduce the SC electroresponsiveness elicited by somatic current injections (Deb et al., 2002; Zitzler and Kunzli, 2004; Druckmann et al., 2007; Druckmann et al., 2008)

The optimization workflow followed the same procedure used for parameters tuning in GrCs (Masoli et al., 2017; Masoli et al., 2020). A set of 36 G_i-max_ values was tuned to match the firing pattern revealed by electrophysiological recordings. From the experimental traces, used as templates, we extracted the experimental features necessary to assess the fitness functions for the optimization procedure.

The following features, selected to better reproduce the biophysical properties of action potentials, were extracted using the ‘Electrophys Feature Extraction Library’ (eFEL) (Van Geit, 2015) during the spontaneous firing and the current injections protocols: action potential amplitude (AP_Ampl_) (mV), action potential afterhyperpolarization (AP_AHP_) (mV), action potential threshold (AP_Thr_) (mV), action potential half-width (AP_HW_) (ms), action potential frequency (Hz) (Supplementary Tables 4 and 5).

Optimization tests were applied to have four different best models based on the four morphologies, starting the process with the same range of conductances. Tests were performed on Piz Daint Cluster (CSCS – Lugano), using 2 nodes for a total of 72 cores, with simulations of 2000 ms, fixed steps of 0.025 ms, temperature of 32 °C, as in the experimental recordings and all ionic current kinetics were normalized to this value using Q_10_ corrections (Gutfreund et al., 1995). The stimulation protocol included the spontaneous firing, two positive current injections (4, 16 pA) and a negative current injection (−16 pA) lasting for 2 s. The optimizations involving 100 individuals for 50 generations, required 3 h of computation time.

#### Model simulations and testing protocols

The best models yielded by the optimization process were simulated using protocols designed to reproduce the experimental ones. Each model was tested for: (i) the spontaneous firing frequency, (ii) the firing frequency obtained with positive current injections of 4 pA and 16 pA, (iii) the AHP after current injection and (iv) the sag generated by −16 pA current injection. The models were further tested with synaptic protocols. (v) PF activation was tested with synaptic trains composed by 10 pulses @ 4 Hz, 10 Hz, 20 Hz, 50 Hz, 100 Hz, 200 Hz, 500 Hz. (vi) GABAergic inhibitory activity, between two pair of SC, was tested using a background stimulation at 20 Hz with either 20, 27 or 32 GABA_A_ receptor-mediated synapses. The same PF trains were delivered in conjunction with the inhibitory background.

The SC model was also wired in a minimal microcircuit including PFs and a PC (Masoli and D’Angelo, 2017). All the synapses involved were randomly distributed over the corresponding neuronal compartments. The PF formed 100 synapses on PC dendrites and 3 synapses on SC dendrites, the SC synapses formed 25, 50, 100, 200 or 300 synapses on PC dendrites. In a set of simulations, a second SC was connected through 32 synapses to the first one. SCs and the PC received PF stimuli made of 10 pulses @ 4 Hz, 10 Hz, 20 Hz, 50 Hz, 100 Hz, 200 Hz or 500 Hz.

The simulations were performed on an AMD Threadripper 1950X (32GB ram), with fixed time step of 0.025 ms, temperature 32°. Simulation time was 2 s for spontaneous activity and current injections, 5 s to test PF-SC synaptic activity, 15 sec to test the SC –> SC –>PC circuit.

### Data analysis

Custom Python-scripts were written to automatically analyze the set of G_i-max_ parameters at the end of each optimization, to run simulations and to extract features. The voltage and current traces were analyzed using MATLAB 2018a/b (MathWorks, Natick, MA, USA), pClamp 10 software (Molecular Devices, CA, USA), OriginPro software and MS Excel. The morphologies were analyzed with NEURON and visualized with Vaa3D.

### Model dissemination

The experimental traces can be found on the HBP knowledge Graph at this link: https://kg.ebrains.eu/search/instances/Dataset/3ca4af33-64bd-437a-9c53-2dac19e10168. The models will be made available on the Brain Simulation Platform (BSP) of the Human Brain Project (HBP) as a “live paper” containing a selection of routines and optimization scripts. A wider selection will be made available on the HBP knowledge Graph. The models will also be uploaded on ModelDB.

## Results

### Stellate cell physiology and modeling

Detailed models of SCs of mouse cerebellum were generated starting from morphological reconstructions and using electrophysiological recordings as a template for the optimization of membrane mechanisms (Masoli et al., 2017; Masoli et al., 2020) (full explanation is given in Supplemental Material). Morphologies of 4 SCs were reconstructed from immunofluorescent images taken from fixed preparations *ex vivo*. WCR from 23 SCs were obtained in acute cerebellar slices. All neurons were selected in the outer 2/3^rds^ of the molecular layer, where SCs are the only neuronal species present (Chan-Palay and Palay, 1972).

The 4 reconstructed SCs corresponded to the canonical description reported in literature (Cajal, 1909,1910; Rakic, 1972; Palay and Chan-Palay, 1974). SCs showed a soma emitting 3-4 branched dendritic trees subdivided in proximal and distal sections, and an initial segment (AIS) continuing in a branched axon (Fig. 1A). The soma surface measured 42.4±9.3 μm^2^ (n = 4). Long, contorted and frequently branching dendrites extending from the soma were characterized centrifugally (Bok, 1959; Uylings et al., 1986), and quantified with two previously established measures (Jacobs et al., 2011; Jacobs et al., 2014): total dendritic length (the summed length of all dendritic segments; 845.2 ± 121.2 μm; n = 4) and segment count (proximal dendrites: 11.0 ±3.0 μm, n = 4; distal dendrites: 63.5 ± 14.0 μm, n = 4). Conversely, the axons branched immediately generating short and circumscribed collaterals (AIS length: 28.0±6.7 μm, n = 4; axon length: 577.9 ± 225.0 μm, n = 4; Supplementary Table 2 and Movie 1). The dendritic tree was flattened on the sagittal plane of the *folium* and the axon, after an initial part parallel to the dendrite, advanced along the transverse plane (Palay and Chan-Palay, 1974; Rieubland et al., 2014; Prestori et al., 2019). The 4 morphologies were transformed into morpho-electrical equivalents (Fig. 1A) and used as the backbone for model reconstruction (Vervaeke et al., 2012; Masoli et al., 2015; Migliore et al., 2018). The SC model was divided into five electrotonic compartments and endowed with 14 different types of voltage-dependent and calcium-dependent ionic channels and with a calcium buffering system (Fig.1B).

**Figure 1.**
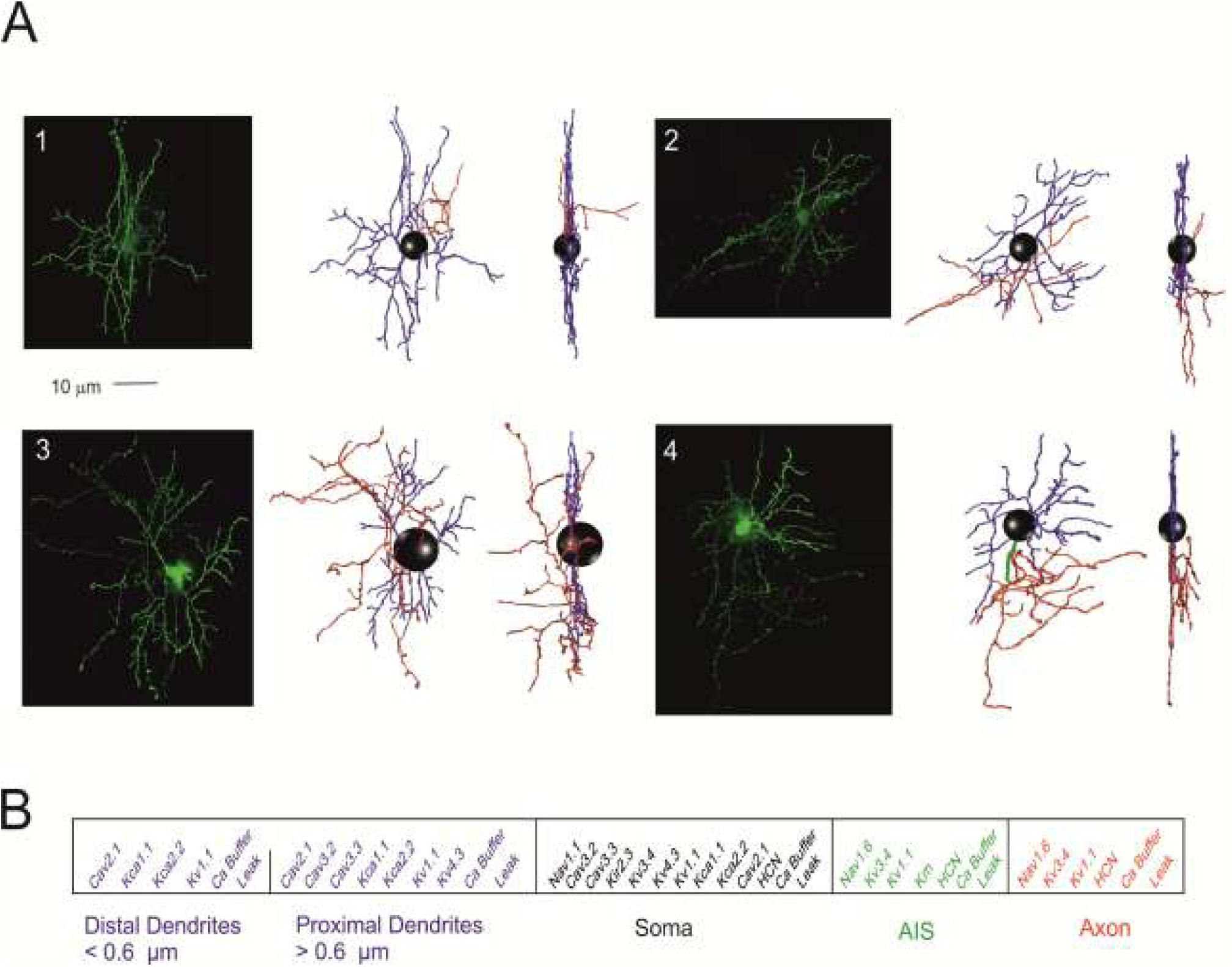
Stellate cell morphological reconstruction. (A) Confocal microscopy imaging of four mouse SCs filled with biocytin (scale bar 10 μm) are shown along with the corresponding digital reconstruction with Neurolucida (visualization with Vaa3D simulator; right). The 3D morphological reconstructions of SCs include dendrites (blue), soma (black), axonal initial segment (green) and axon (red). (B) The SC model is divided into five electrotonic compartments and endowed with specific ionic mechanisms according to immunohistochemical data. Ionic channels include Na^+^, K^+^ and Ca^2+^ channels and a Ca^2+^ buffering system.

The electrophysiological properties of SCs were recorded from neuron located in the outer 2/3^rds^ of the molecular layer (Figs 2–4). The basic properties of SC were homogeneous, according to anatomical indications (Palay and ChanPalay, 1974; see also above), and could be summarized as follows. (i) SC input resistance was 1.3 ± 0.1 GΩ; n = 23 and membrane capacitance was 6.7 ± 0.5 pF; n = 23. (ii) SC spike were fast (1.0 ± 0.2 ms, n = 9; half-width) and overshooting (37.3 ± 5.2 mV) (detailed parameters are reported Supplementary Tables 4 and 5). (iii) SCs fired spontaneous action potentials at 16-36 Hz (24.2 ± 2.1 Hz; n = 9; Fig. 2A). The average ISI distribution showed a single peak at 43.4 ± 3.8 ms (n = 9; Fig. 2B). There was no correlation between spike amplitude and threshold (Fig. 2C).

**Figure 2.**
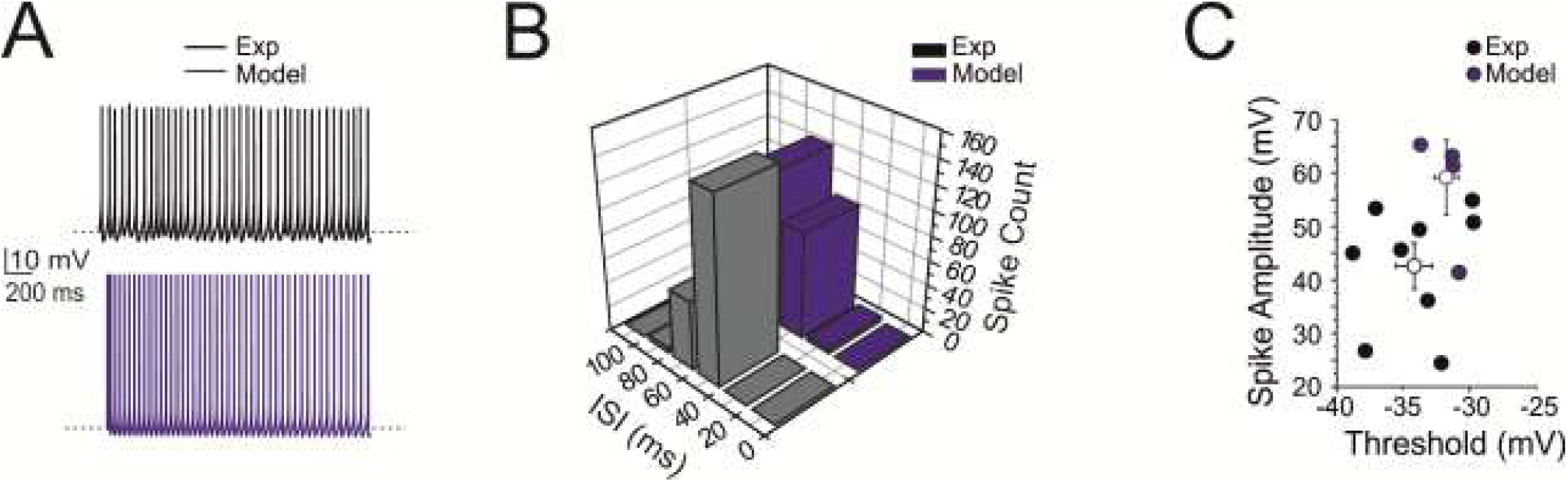
Pacemaking activity. (A) Pacemaker activity during stellate cell WCR (n = 9; black) and in the model (n = 4; blue). (B) Distribution of the ISI of spontaneously firing stellate cells in WCR. Note that the ISI in the model (blue bar; n = 4) falls within the experimental data distribution (black bar; n = 9, p = 0.8). (C) Relationships among ISI parameters. Note that the model data points (blue circles; n = 4) fall within the distribution of spike amplitude vs. spike threshold measured experimentally (black circles; n = 9). The model did not significantly differ from the experimental data (p = 0.64). Data are reported as mean ± SEM.

(iv) When injected with hyperpolarizing current steps, SCs showed sagging inward rectification and, after hyperpolarization, showed rebound excitation (Fig. 3A). Rebound excitation consisted of accelerated spike frequency, which progressively decayed back to baseline. There was no correlation between the amplitude of the hyperpolarizing sag (10.3 ± 2.0 mV; n = 5), the first spike delay (17.2 ± 2.3 ms; n = 5) or the first ISI (20.1 ± 1.0 ms; n = 5) (Fig. 3B).

**Figure 3.**
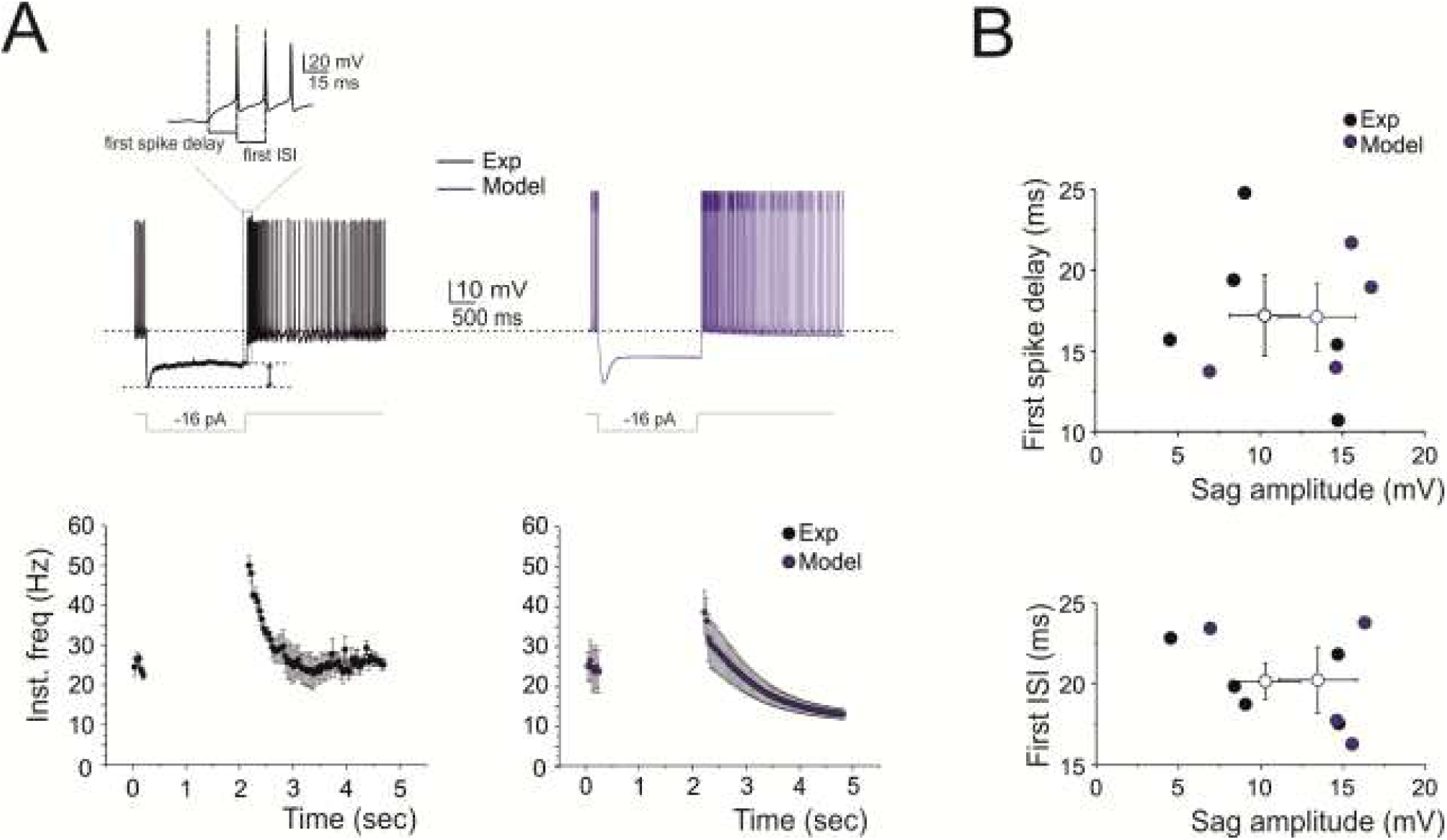
Response to hyperpolarization and rebound excitation. (A) A WCR from a SC shows sagging inward rectification in response to hyperpolarizing current injection and, at the end of the hyperpolarization, rebound excitation with an early and a protracted phase of intensified firing. Simulations of this specific experiment show that the model could faithfully reproduce sagging inward rectification and rebound excitation (blue trace). At the bottom, the time course of spike frequency for a −16 pA current pulse. (B) The plots report the time to first spike and the first ISI during rebound excitation as a function of sag amplitude (black circles; n = 5). Note that the model (blue circles; n = 4) did not significantly differ from the experimental data (p = 0.32). Data are reported as mean ± SEM.

(v) When injected with depolarizing current steps, SCs generated fast repetitive spike discharge. The output frequency increased almost linearly with current intensity (slope 2.79 Hz/pA; n = 5) (Fig. 4A, B). (vi) At the end of spike discharges elicited by step current injections, SCs showed a marked AHP delaying the re-establishment of pacemaker activity. The AHP duration increased with stimulation intensity (Fig. 4B).

**Figure 4.**
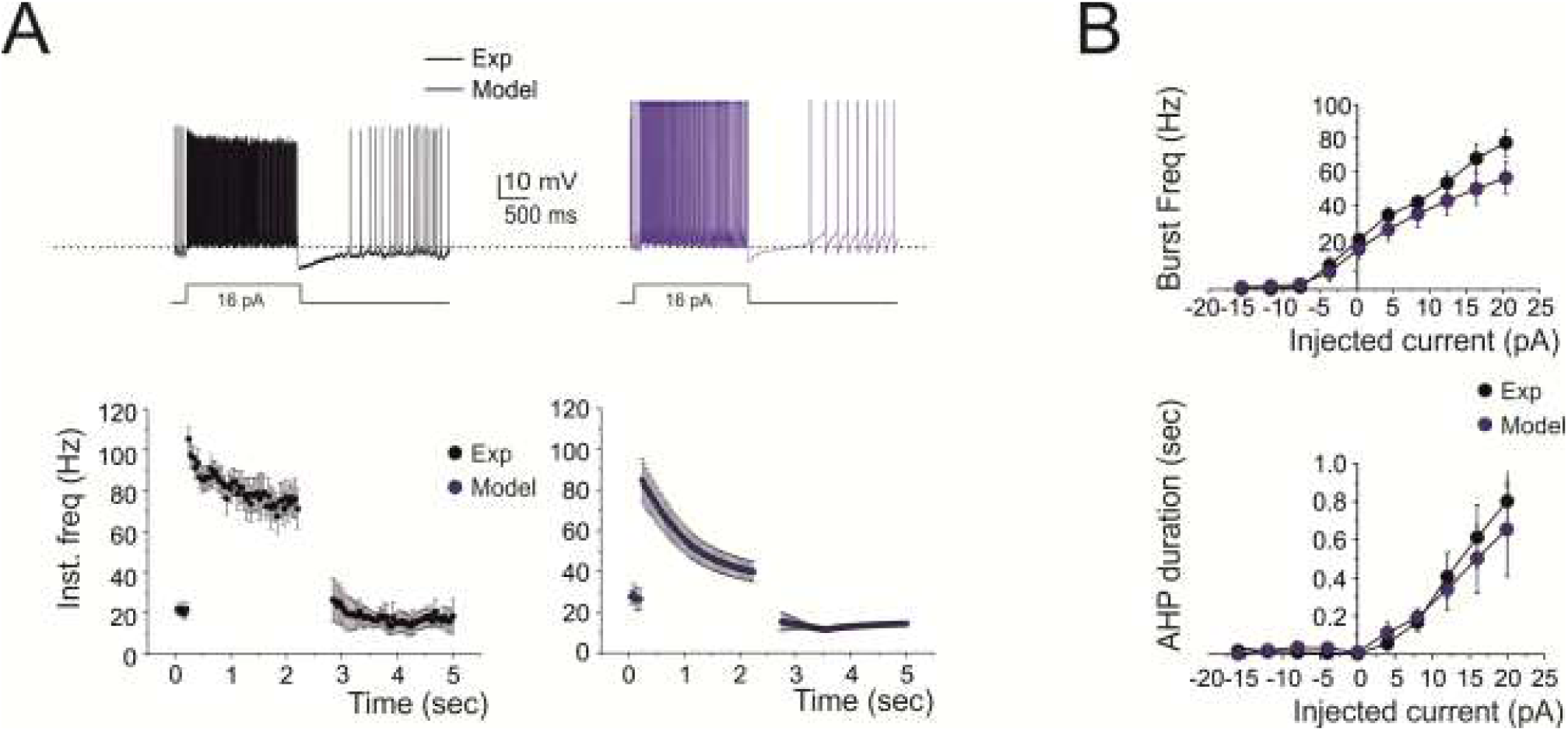
Response to depolarization. (A) A WCR from a SC shows the firing frequency and the pause following depolarizing current step (16 pA). The time course of instantaneous spike frequency for the 16 pA current pulse is shown below. Simulations show that the model could faithfully reproduce this behavior (blue traces). (B) In the plots, the response of the model (blue circles; n = 4) to current injections (from −16 to 20 pA) is compared to experimental data (black circles; n = 5). Simulations show that the model could appropriately fit the experimental measurements of spike frequency versus injected current (16 pA: p = 0.097) and of pause versus injected current (16 pA: p = 0.69). Data are reported as mean ± SEM.

The models (see Methods and Supplemental Material) faithfully reproduced all the properties reported above, as shown in Figures 2, 3 and 4. The statistical comparisons reported below demonstrate the absence of significant differences between the data obtained in WCR and the SC models. In detail, the model (i) matched the passive neuron properties, (ii) reproduced spike shape (Supplementary Tables 4 and 5), (iii) showed an average pacemaker frequency within the experimental data distribution (Figs. 2A, B; Supplementary Movie 2), (iv) responded to negative current injections, reproducing sagging inward rectification and rebound excitation (Fig. 3A), (v) increased firing rate with an almost linear I/O relationship in response to step current injections (slope 2.13 Hz/pA; n = 4) (Fig. 4B), (vi) showed a hyperpolarization followed by a pause in spike firing at the end of the current steps (Fig. 4B). The parameters of the 4 SC models corresponding to these electrophysiological properties fell within the experimental data distributions (p>0.1, unpaired *t*-test), indicating that SCs electrical responses recorded in experiments and models could be treated as members of the same statistical distribution. The electroresponsive mechanisms of the models are fully described in Supplementary Figs. 1 and 3, along with the specific involvement of HCN1, Cav3.2, KCa2.2, Kv4.3 and other voltage-dependent ionic channels reported in SCs.

### Frequency-dependent short-term dynamics at parallel fiber – stellate cell synapses

In order to understand how information transmitted by GrC inputs is encoded by SCs, we investigated response dynamics at PF-SC synapses activated by trains of 20 PF stimuli at different frequencies (50, 100 and 200 Hz) (to preserve PFs, these experiments were performed in coronal slices, see Methods). During the stimulus trains, EPSCs showed an initial strong facilitation followed by depression (Fig. 5A, B) (Bao et al., 2010; Grangeray-Vilmint et al., 2018; Dorgans et al., 2019). The sequence of short-term facilitation and depression was the more evident the higher the stimulation frequency, suggesting that frequency-dependent short-term synaptic dynamics could regulate transmission efficacy along the PF - SC pathway (Mapelli et al., 2010).

**Figure 5.**
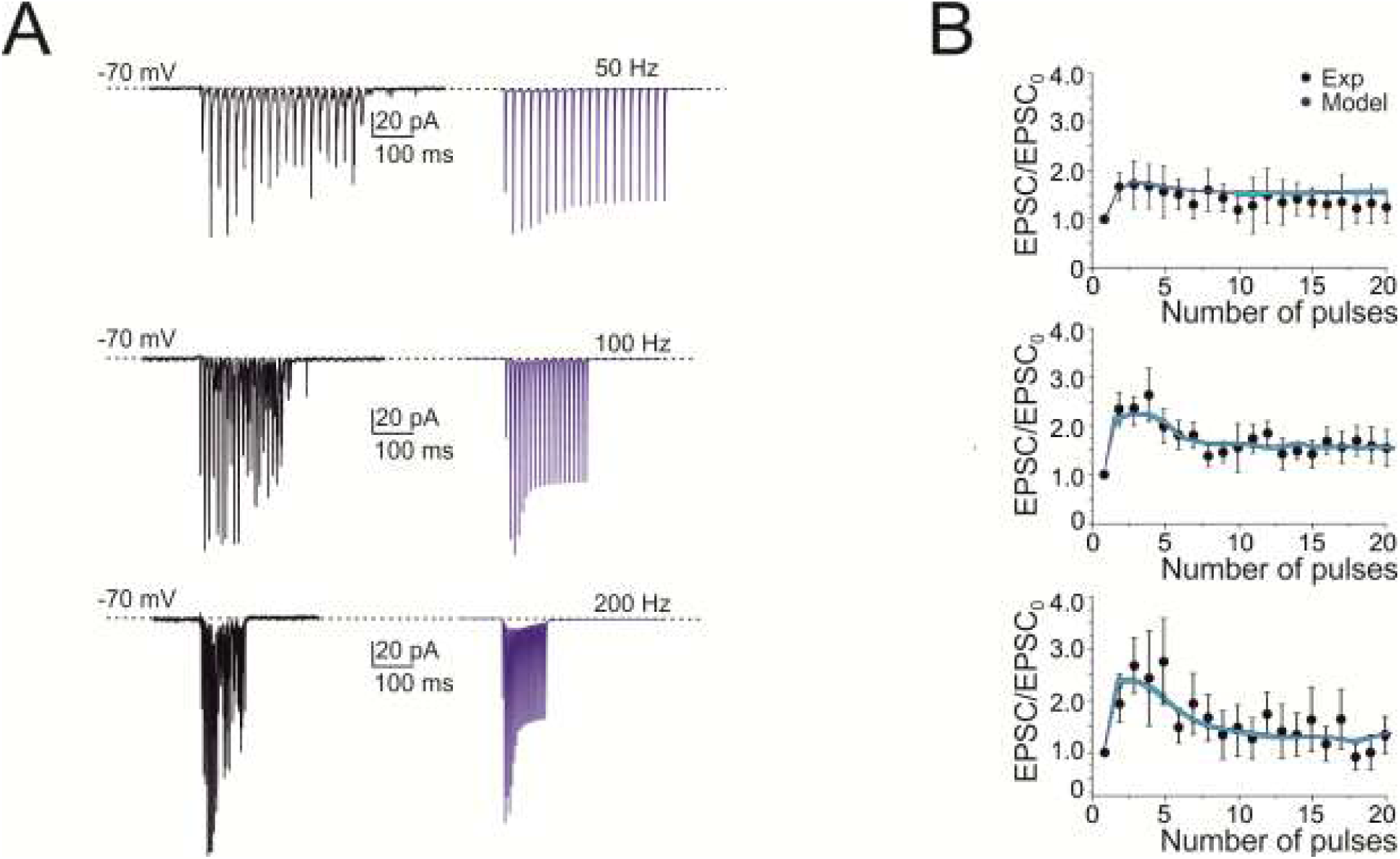
Short-term plasticity at PF-SC synapses. (A) EPSCs recorded from the SC during 50, 100 and 200 Hz parallel fiber stimulations (n = 6). Cells were voltage-clamped at −70 mV and in the presence of gabazine. Each trace (black) is an average of 10 sweeps. Note that EPSC trains showed first facilitation and then depression. Simulations show that the model could faithfully reproduce the same behavior (blue traces; n = 4). (B) Comparison of peak amplitudes of experimental and simulated EPSCs during high-frequency parallel fiber stimulation. Amplitudes are normalized to the first EPSC in the train (correlation coefficient R^2^ @50 Hz = 0.80; @100 Hz = 0.90; @200 Hz = 0.84). Data are reported as mean ± SEM.

The PF - SC model of synaptic transmission (Tsodyks et al., 1998; Nieus et al., 2006) was tuned to follow the experimental results yielding release probability, *p* = 0.15 (Carter and Regehr, 2000; Nieus et al., 2006; Bidoret et al., 2015). The low *p* level generated short-term facilitation, while vesicle depletion during the train caused the subsequent short-term depression (Fig. 6A). The model precisely followed the experimental results (cf. Fig. 5B). The NMDA receptors enhanced PF-SC synaptic transmission (Fig. 6A inset; see also Fig. 8B).

**Figure 6.**
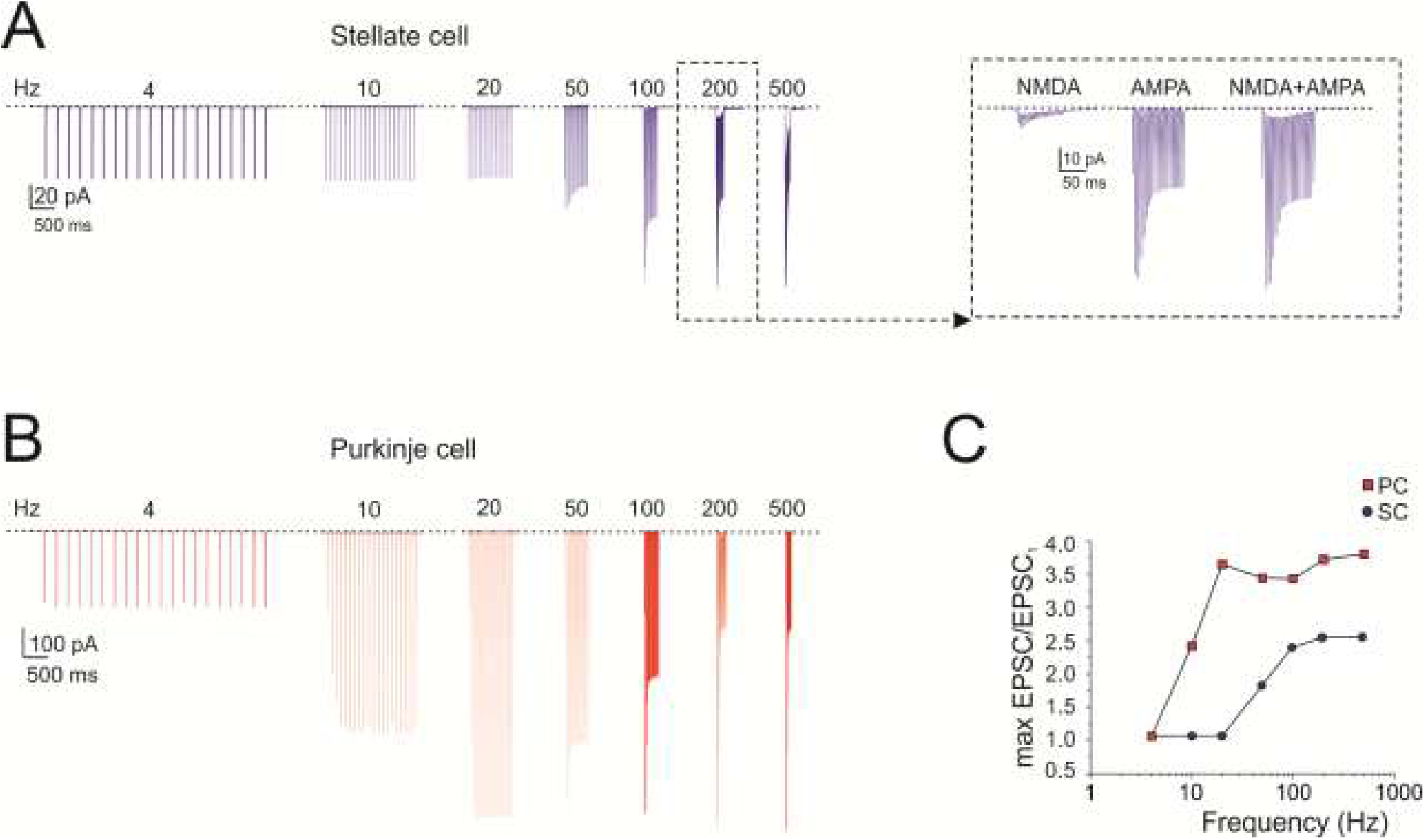
Burst currents at PF-SC and PF-PC cell synapses. (A) The traces show simulated SC EPSCs during trains of 20 stimuli delivered to PFs at different frequencies (4, 10, 20, 50, 100, 200 and 500 Hz; n = 4). Inset, NMDA, AMPA and NMDA+AMPA currents at 200Hz. (B) The traces show simulated PC EPSCs during trains of 20 stimuli delivered to PFs at different frequencies (4, 10, 20, 50, 100, 200 and 500 Hz). (C) Gain curve for SC (blue circles; n = 4 for each frequency) and PC (red squares) responses with respect to input burst frequency. Gain is the ratio between the maximum response obtained at a certain frequency and the first EPSC. The gain curves show a sigmoidal shape with 50% amplitude around 50 Hz for the SC and around 10 Hz for the PC. Data in C corresponds to the cells in A and B.

For comparison, we have run a similar simulation on a PC model (Masoli et al., 2015; Masoli and D’Angelo, 2017), resulting in a similar sequence of short-term facilitation and depression but with different frequency-dependence (Fig. 6B). This difference suggested that the two cells, if wired together, would generate specific filtering effects (see below). The gain curve with respect to input burst frequency showed a sigmoidal shape with 50% amplitude around 50 Hz for the SC and around 10 Hz for the PC (Fig. 6C).

### Frequency-dependence of stellate cell input-output gain functions

The impact of short-term dynamics at PF-SC synapses was investigate by measuring the responses to PF bursts at different frequencies (10 pulses @ 4 Hz, 10 Hz, 20 Hz, 50 Hz, 100 Hz, 200 Hz, 500 Hz). Pronounced spike burst were generated at high frequencies (e.g. at 100 Hz, the frequency increase was 154.5±40.5%, n = 4; p = 0.046; Fig. 7A). A careful analysis of PSTHs showed a delay in the first spikes of SCs in response to high-frequency PF stimulation (about 50 ms at 100 Hz) probably determined by short-term facilitation in synaptic transmission (Fig. 7B). The analysis of responses to different PF input frequencies revealed that SC responses did not increase their spike output frequency above the basal frequency below about 10 Hz, then their responses increased and tended to saturate beyond 100 Hz (Fig. 7C). The input/output gain curve showed a sigmoidal shape (Fig. 7D).

**Figure 7.**
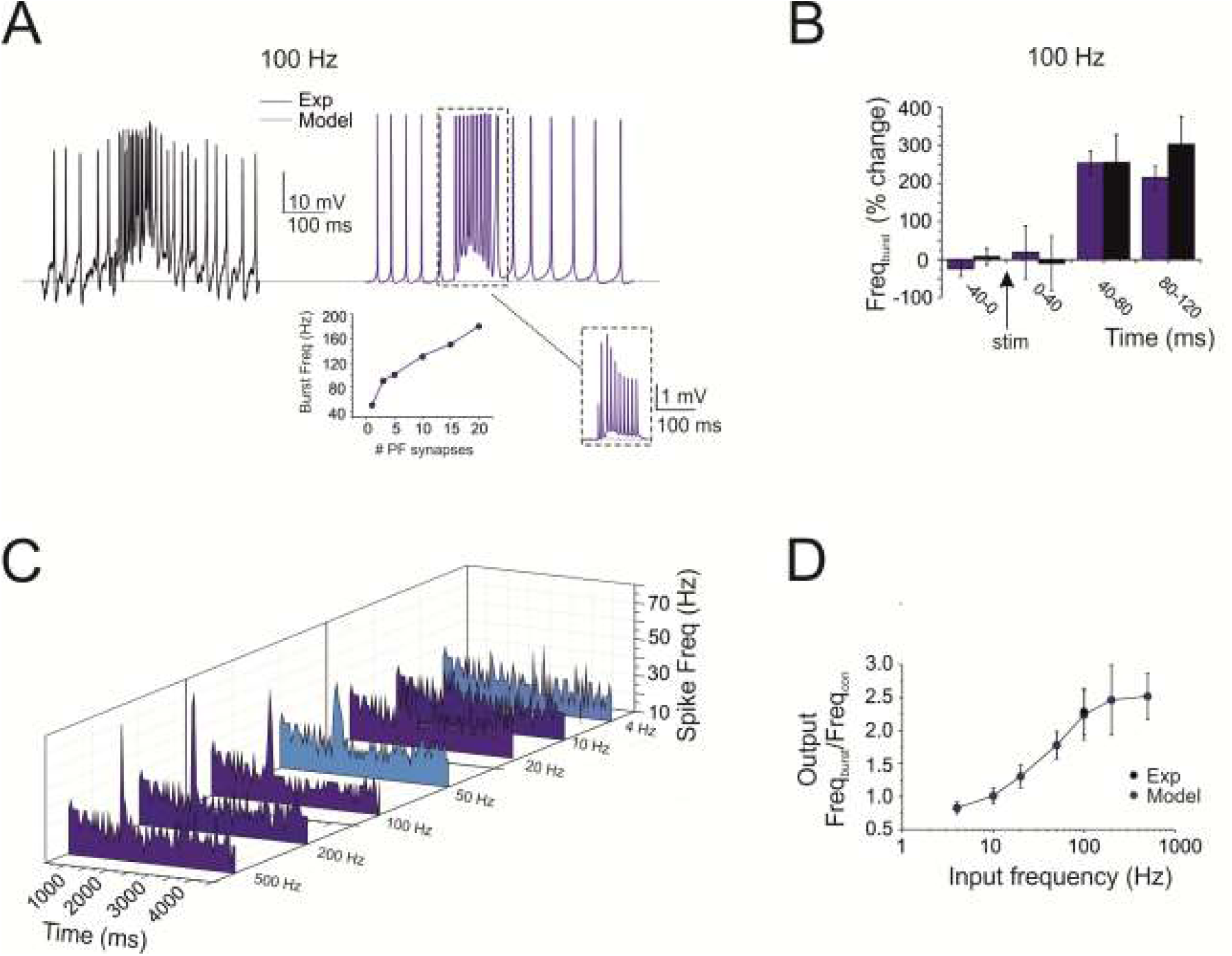
Frequency-dependence of SC input-output gain function. (A) The traces show a SC burst in response to 10 pulses @ 100 Hz-delivered to PFs (black trace) and the corresponding simulation (blue trace). Inset (left), calibration of the number of PF-SC synapses required to obtain a given burst frequency (3 in the example). Inset (right), the synaptic current (AMPA+NMDA) corresponding to the simulated burst. (B) The histogram shows the time course of the SC burst shown in A in response to 10 pulses @ 100 Hz-delivered to PFs (black trace) and the corresponding simulation (blue trace). Note the about 50 ms delay to burst response. (C) The array of PSTHs shows the SC responses to PF bursts at different frequencies (10 pulses @ 4 Hz, 10 Hz, 20 Hz, 50 Hz, 100 Hz, 200 Hz, 500 Hz). Note that pronounced spike bursts were generated at high frequencies (e.g at 100 Hz). (D) Input/output SC gain for experimental (black, n = 5) and simulated bursts (blue, n = 4). SCs did not increase their spike output frequency until about 10 Hz, then their responses increased and tended to saturate beyond 100 Hz. Note the superposition of the single experimental data point and simulated data. Data in B and D are reported as mean ± SEM (n = 4).

The models were calibrated by activating an increasing number of synapses at 100 Hz, reveling that an output frequency like the experimental one was obtained using just 3 PF-SC synapses (Fig. 7A, inset). The experimental data were then faithfully reproduced by the models (Fig. 7 A-D; see also Supplementary Figs. 2) by simply using the electroresponsiveness and synaptic transmission properties calibrated beforehand, thus providing a mechanistic explanation for the time-dependent and frequency-dependent properties of SCs synaptic responsiveness.

The model was further used to evaluate the effect of factors that could modulate the gain curve. NMDA current stimulation ((Bower and Woolston, 1983; Carter and Regehr, 2000; Clark and Cull-Candy, 2002; Nahir and Jahr, 2013) blockage reduced (though not significantly) the SC firing rate during high-frequency bursts (e.g. at 200 Hz, frequency change was −21.4 ± 9.2%, n = 4; p = 0.13; Fig. 8A, B). Of special interest was the simulated effect of inhibitory GABA_A_ receptor-mediated synaptic transmission from neighboring SCs (Kondo and Marty, 1998; Pouzat and Marty, 1999; Alcami and Marty, 2013; Nieus et al., 2014). To estimate the strength of this connectivity, a series of simulation were performed with different numbers of inhibitory synapses activated simultaneously with PF bursts. While increasing the inhibitory strength, SC excitation was moved toward higher frequencies and the activation curve became steeper (50 Hz _(mod)_: −70.9 ± 9.4%, n = 4; p = 0.009; Fig. 8B). The activation of voltage-dependent currents in the dendrites by delivering multisynaptic stimulation pattern to the SC model is fully described in Supplemental Material (Supplementary Fig. 4). In conclusion, during PF stimulation, NMDA receptor modulation was more evident at higher frequencies whereas GABAergic modulation was more evident at lower frequencies suggesting that two synaptic systems had opposite effects on the SC gain function.

**Figure 8.**
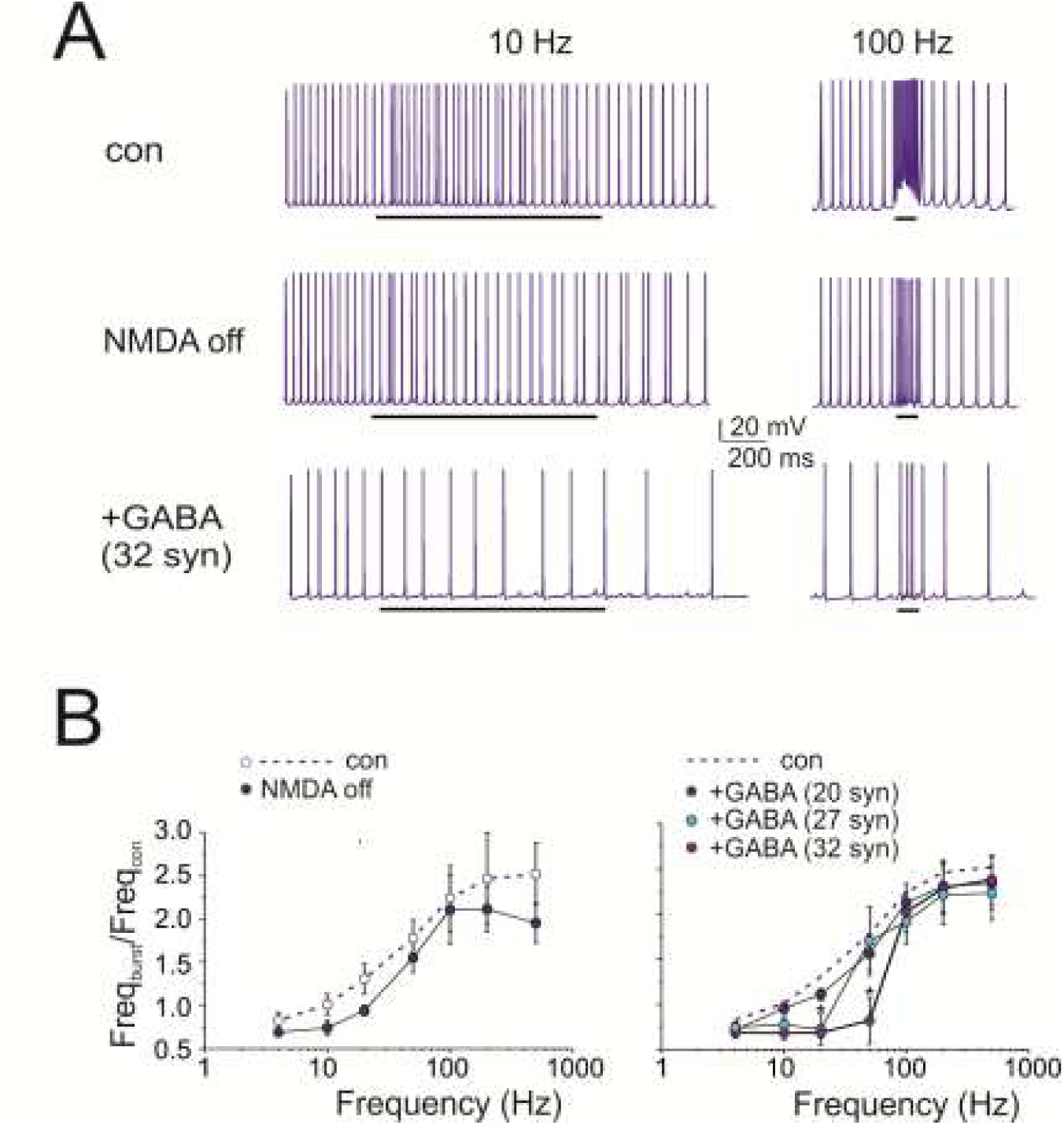
Regulation of SC gain function by NMDA and GABA_A_ receptors. (A) The traces show simulated SC response to 10 pulses @ 100 Hz-delivered to PFs. NMDA receptor switch-off and activation of inhibitory (32) synapses activated simultaneously with the PF burst reduced SC firing rate. Black bar indicates the stimulus duration. (B) Input/output SC gain regulation. Note that the NMDA current block reduced the spike output frequency mostly during high-frequency bursts (200-500 Hz), while a sufficient number of inhibitory synapses (activated simultaneously with PF bursts) shifted SC excitation toward higher input frequencies. Data are reported as mean ± SEM (n = 4).

### Prediction of stellate cell filtering of Purkinje cell responses along parallel fiber beams

SCs are known to provide feed-forward inhibition to PCs along PF beams (Santamaria et al., 2002; Santamaria and Bower, 2005; Santamaria et al., 2007; Bower, 2010; Masoli and D’Angelo, 2017). Here, the different frequency-dependence of SC and PC responses to input bursts suggested that SCs could act as filters, when the two cells are co-activated along a PF beam (Fig. 9A).

**Figure 9.**
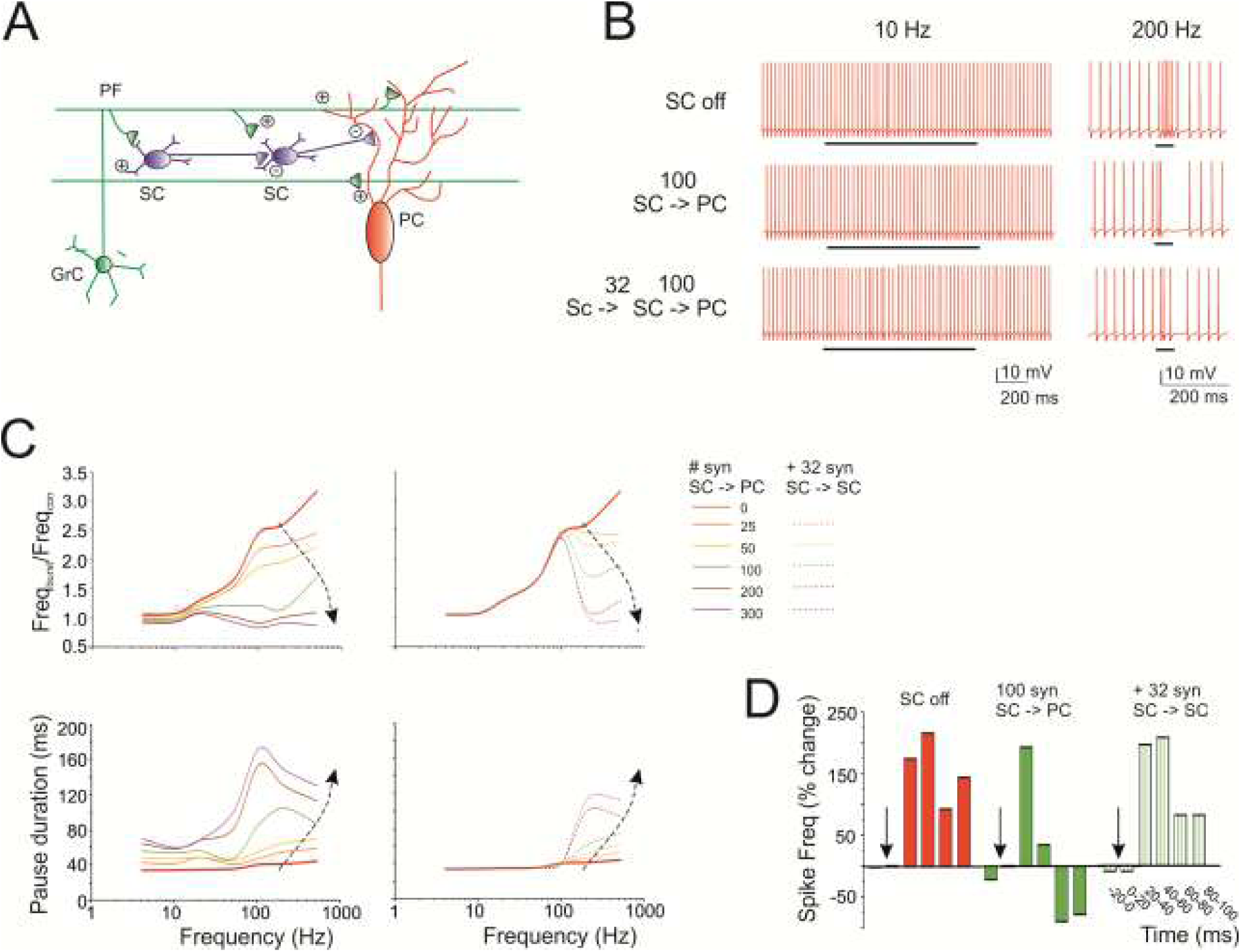
Prediction of SC filtering of PC responses along PF beams. (A) Schematics of the afferent connections to a PC activated by PF stimulation. Granule cell (GrC); parallel fiber (PF); stellate cell (SC); Purkinje cell (PC). The figure highlights the interactions of elements in the cerebellar molecular layer and the location of afferent PC synapses. (B) The traces show simulated PC response to 10 pulses at different frequencies (10, 200 and 500 Hz) delivered to 100 PFs in which (i) SCs were not activated (SC off), (ii) 100 SC synapses were activated (SC –> PC) and (iii) 100 SC synapses received inhibition from 32 SC synapses (SC –> SC –> PC). Black bar indicates the stimulus duration. (C) Input/output PC burst frequency gain (left) and pause length (right). Different curves are obtained using PF trains at different frequency (10 pulses @ 4 Hz, 10 Hz, 20 Hz, 50 Hz, 100 Hz, 200 Hz, 500 Hz) and an increasing number of inhibitory synapses. Dotted traces also include the case of SC-SC inhibition. Note that PC burst and pause showed an almost opposite modulation by SCs. (D) The histogram shows the regulation of PC firing frequency when SCs are off, on and reciprocally inhibited. Note that reciprocal SC inhibition can abolish the effect of SCs on PCs.

To explore the issue, we performed simulations in which SCs were connected to PCs “on beam” (Supplementary Movie 4, 5). When SC synapses were activated, the PC was inhibited and its output spike frequency decreased. The effect was stronger at high input frequencies, where SCs are more activated. Consequently, a sufficient number of SC synapses (>=50 Hz) abolished the response of PCs at high frequency (>50 Hz) without affecting that at low frequency (<50 Hz), *de facto* flattening the gain curve. In this configuration, the SC – PC system acted as an effective low-pass filter (Fig. 9 B, C).

SCs are wired together through inhibitory synapses. We therefore performed simulations in which, in addition to stellate-PC feed-forward inhibition, SCs were reciprocally inhibited. In this case, the effect on the PC gain curve changed. Reciprocal inhibition reduced SC activity above 50 Hz, while leaving the PC excited above 10 Hz (cf. Fig 6C). In this configuration, the SC –> SC –> PC system acted as low-pass filter was partially deactivated (Fig. 9 B, C).

PCs normally generate burst-pause responses following PF bursts (Herzfeld et al., 2015; Masoli and D’Angelo, 2017). The simulations showed that, along with the burst, the SCs also regulated the pause although in an opposite manner (Fig. 9 B, C). The pause was almost insensitive to input frequency but increased remarkably along with SC inhibition. Finally, SCs activity delayed PC responses by about 20 ms (Fig. 9D). In aggregate, these simulations show that SCs exert a remarkable effect on the delay, band-pass and burst-pause behavior of PCs.

## Discussion

This work provides the first detailed models of a small set of cerebellar SCs. The models are based on precise morphological and electrophysiological measurements and faithfully reproduce the non-linear excitable properties and short-term neurotransmission dynamics at the PF - SC synapse. The models give insight on the critical role that these neurons could play in molecular layer processing. By exploiting membrane excitability and short-term synaptic plasticity, SCs recode PF bursts and regulate the gain of PCs in a frequency-dependent manner. SC inhibition of PCs occurs above 10 Hz and, depending on the engagement of SC-SC inhibitory chains, can generate low-pass or band pass filters. These observations expand the concept of the cerebellum as an adaptive filter (Dean and Porrill, 2010).

### Stellate cell electroresponsiveness and synaptic regulation

SCs showed autorhythmic firing around 25 Hz (Hausser and Clark, 1997; Carter and Regehr, 2002). In slice preparations, PFs and climbing fibers are inactive (D’Angelo et al., 1995), and the only connection conveying spontaneous activity comes from other SCs. It is thus probable that SC pacing reflected intrinsic electroresponsive properties since, in our recordings, GABAergic inputs were blocked pharmacologically. When stimulated with positive or negative currents, SCs showed burst-pause or pause-burst responses resembling those of other cerebellar neurons, comprising PCs (De Zeeuw et al., 2011; Masoli and D’Angelo, 2017), Golgi cells (Forti et al., 2006; Solinas et al., 2007b, a) and deep nuclear cells (Dykstra et al., 2016; Moscato et al., 2019). Therefore, SCs are well suited to take part into the complex bursting and pausing dynamics of the cerebellar circuit (Herzfeld et al., 2015).

While several ionic channels have been reported in these neurons, the models provides insight on how they operate during SC activity (see Supplemental Material for details). LVA channels (Cav3.2 and Cav3.3) (Molineux et al., 2005; Molineux et al., 2006; Anderson et al., 2013), H channels (HCN1) (Luján et al., 2005; Angelo et al., 2007) and KCa channels (KCa2.1 KCa2.2) (Womack et al., 2009; Kaufmann et al., 2010; Rehak et al., 2013; Turner and Zamponi, 2014) were critical in driving bursts, pauses and rebounds, while faster channels controlled action potential generation and repolarization. The A-channel (Kv4.3) regulated first spike delay in rebound bursts. The same ionic currents were activated during repetitive synaptic transmission.

Synaptic currents reflected the specific receptor-channel properties reported in voltage-clamp recordings (Carter and Regehr, 2000; Nieus et al., 2006; Nieus et al., 2014; Bidoret et al., 2015) and, when coupled to a presynaptic release mechanism, faithfully accounted for repetitive neurotransmission in bursts at different frequencies. While AMPA receptor-mediated currents generated rapid EPSCs and fast membrane depolarization, NMDA currents caused a slower build-up of the response during the trains. GABA_A_ receptors generated SC inhibitory currents. Interestingly, the kinetics of synaptic response trains showed strong facilitation (short-term depression followed the first impulses at >50 Hz) that, in the model, implied a low neurotransmitter release probability (*p* = 0.15). Consequently, the SCs generated spikes with a delay (about 50 ms during a 100 Hz train) and prevented responses to single PF spikes. This mechanism therefore effectively controls the timing of SC responses to bursts at the same time preventing their activation by single sparse PF spikes, which are therefore filtered out as noise.

As long as the models predict that SCs operate as delayed high-pass filters, they also anticipate the underlying mechanisms. The SC gain curves were enhanced by NMDA and reduced by GABA_A_ receptor-mediated transmission, this latter also shifting the cut-off frequency from ~50 Hz to ~100 Hz. The frequency-dependent effect reflected the short-term synaptic dynamics typical of these neurons and were enhanced by activation of NMDA currents and voltage-dependent ionic currents (mostly the LVA calcium current). GABA_A_ currents reduced the cell input resistance and prevented regenerative depolarizing effects (see also Supplemental Material).

### Stellate cell regulation of Purkinje cell gain

As well as SCs, also PCs showed short-term facilitation and their gain increased with input frequency (cut-off ~20 Hz). When the SC and PC models were put in series along the PF beam so as to emulate a feed-forward inhibitory circuit (Santamaria et al., 2002; Santamaria and Bower, 2005; Santamaria et al., 2007; Bower, 2010; Masoli and D’Angelo, 2017), the gain of PCs was limited to the values typical of low-frequency transmission (4-10 Hz) generating a low-pass filter. When the SC–SC feed-forward inhibition (Mittmann et al., 2005; Rieubland et al., 2014) was activated, the gain curve of the PC showed a decrease beyond 100 Hz, generating a band-pass filter.

Validation of these simulations is provided by experiments that were performed using voltage-sensitive dyes in coronal slices. While PCs on-beam showed increased gain at high frequency when GABA_A_ receptors were blocked, their responses were low-pass filtered to the 4-10 Hz range when synaptic inhibition was active (Mapelli et al., 2010). In addition to this, spots of PCs receiving activation along vertical beams, proved capable of high-frequency gain probably reflecting the exceeding number of GrC ascending axon synapses on PCs compared to the low number of PF synapses on local SCs (Bower and Woolston, 1983; Cohen and Yarom, 2000). Thus, while evidence for low-pass filtering is available, that for band-pass filtering awaits for experimental validation.

### The molecular layer as an adaptive filter

A main prediction of th present simulations is that the intensity and bandwidth of molecular layer filtering is modulated by the number of active synapses between PFs, SCs and PCs. The number of active synapses, in turn, would depend on the organization of spike discharge in the PFs, and therefore, ultimately, on the reaction of the granular layer to the mossy fibers input (Gall et al., 2005; D’Angelo and De Zeeuw, 2009; D’Errico et al., 2009). Moreover, in addition to PF - PC synapses (Ito and Kano, 1982; Salin et al., 1996; Lev-Ram et al., 2002; Coesmans et al., 2004; Qiu and Knöpfel, 2009), also the synapses between PFs and SCs (Rancillac and Crépel, 2004; Liu et al., 2008; Bender et al., 2009) and between SCs and PCs (Kano et al., 1992; Kawaguchi and Hirano, 2007; Hirano and Kawaguchi, 2014) have been reported to develop long-term synaptic plasticity [for a recent review see (Gao et al., 2012; D’Angelo, 2014; Mapelli et al., 2015)]. It is therefore possible that the PF-SC-PC system operates as a bank of filters, in which gain and bandwidth of PC responses are finetuned by SCs. Interestingly, mechanisms of signal filtering tuned by local synaptic plasticity have recently been recognized also in the cerebellum granular layer (Casali et al., 2020; Masoli et al., 2020) extending the mechanisms that could actually transform the cerebellum into an adaptive filter (Dean and Porrill, 2010).

## Conclusions

In conclusion, SCs emerge as critical elements controlling cerebellar processing in the time and frequency domain. In combination with VSD recordings (Mapelli et al., 2010), the present simulations support the view that computation along PF beams is different from that occurring in the spots of PCs laying vertically on top of active granular layer clusters (Bower and Woolston, 1983; Bower, 2002; Diwakar et al., 2011). Tuning of transmission bandwidth and delay through specific membrane and synaptic mechanisms may contribute to explain the role of SCs in motor learning and control (Jorntell et al., 2010). For example, transmission of low-frequency bursts along the beams may help tuning the molecular layer with cortical low-frequency oscillatory inputs on the theta-band (Ros et al., 2009; Courtemanche et al., 2013), with reflections on circuit synchronization and on the induction of long-term synaptic plasticity. These potential consequences remain to be investigated using advanced recording techniques and large-scale circuit simulations (Casali et al., 2019; Arlt and Häusser, 2020).

## Author contribution

MR has fully developed the SC models and the corresponding simulation analysis and text. FL performed the patch-clamp recordings. SM supervised modeling and simulations with Purkinje cells. DSP and AM performed the morphological SC reconstructions. FP supervised experiments \and data analysis and wrote the first version of the paper. ED coordinated the whole work, participated to data analysis and model simulations and wrote the final version of the paper.

