## Supplemental Material for "Stellate cell computational modelling predicts signal filtering in the molecular layer circuit of cerebellum"

#### *SUPPLEMENTARY NOTES*

List of abbreviations used below:

- SC, stellate cell
- PC, Purkinje cell
- AIS, axon initial segment
- Dendprox, proximal dendrites
- Denddist, distal dendrites
- AP, action potential
- AHP, afterhyperpolarization
- Thr, threshold
- Ampl, amplitude
- HW, half-width
- Freq, frequency
- eFEL, Electrophys Feature Extraction Library

##### *Ionic channels*

Nav1.1 - Nav1.6. Expression and distribution of SC sodium channels were determined experimentally (Schaller and Caldwell, 2003; Lorincz and Nusser, 2008). Nav1.1 channel was placed on the soma and Nav1.6 on the AIS and axonal compartment. The gating mechanism was taken from (Khaliq et al., 2003; Magistretti et al., 2006).

Kv1.1. The low threshold Kv1.1 channel, in accordance with experiments (Southan and Robertson, 2000; Lorincz and Nusser, 2008; Williams et al., 2012), was placed on all compartments. The gating mechanism was taken from (Akemann and Knöpfel, 2006).

Kv3.4. This ionic channel with delayed rectifier properties was distributed on the soma, AIS and axon to repolarize the $Na^{+}$ spikes (Perney et al., 1992; Brooke et al., 2006; Rowan et al., 2014). The gating mechanism was taken from (Akemann and Knöpfel, 2006).

Kv4.3. This ionic channel with A-type properties was placed on the soma and proximal dendrites (Molineux et al., 2005; Anderson et al., 2013). Moreover, Kv4.3 interacts with the LVA $Ca^{2+}$ (Cav3.x) channels to create a complex with important functions in SC firing (Turner and Zamponi, 2014). The gating mechanism was taken from (Masoli et al., 2015).

Kv7.x. The M-current was identified electrophysiologically (Pan et al., 2006; Brown and Passmore, 2009; Miceli et al., 2012). Kv7 channels were expressed in the SC AIS were placed in the AIS using gating mechanisms developed for the granule cell (D'Angelo et al., 2001).

Kir. Inward rectifier $K^+$ channel was expressed in the SC soma (Stonehouse et al., 1999; Prüss et al., 2005; Raphemot et al., 2011).

KCa1.1 - KCa2.2. Large and small conductance calcium-activated potassium channels, which can cluster with Cav2.1 channels, were placed on the proximal/distal dendritic and somatic compartments based on immunohistochemical and electrophysiological data (Womack et al., 2009; Kaufmann et al., 2010; Rehak et al., 2013; Turner and Zamponi, 2014). The gating mechanism was taken from (Anwar et al., 2012).

Cav2.1. The high-threshold calcium channels (P-type) were placed on the proximal/distal dendritic and somatic compartments (Kulik et al., 2004; Indriati et al., 2013). The gating mechanism of Cav2.1 was taken from (Swensen and Bean, 2003; Anwar et al., 2012).

Cav3.2 – Cav3.3. The low-threshold calcium channels (T-type) were placed on the proximal dendritic and somatic compartments (Molineux et al., 2005; Molineux et al., 2006; Cain and Snutch, 2010; Anderson et al., 2013; Turner and Zamponi, 2014). The gating mechanism of Cav2.1 was taken from (Huguenard and McCormick, 1992; Xu and Clancy, 2008).

HCN1. Hyperpolarization activated cyclic nucleotide-gated cationic channel) was placed on the AIS, somatic and axonal compartments (Luján et al., 2005; Angelo et al., 2007). The gating mechanism was taken from (Solinas et al., 2007a, b).

Calcium dynamics. The calcium buffer was taken from (Masoli et al., 2015) and modified to contain Parvalbumin, the typical calcium binding proteins of the SC (Bastianelli, 2003; Collin et al., 2005; Anwar et al., 2012; Alcami and Marty, 2013).

*SUPPLEMENTARY TABLES*

#### *Supplementary Table 1. Ionic mechanisms in stellate cell models*

| Ionic Channel | Location | Range Gli-max (mS/cm$^2$) | $E_{rev}$ (mV) |
|---|---|---|---|
| Nav1.1 | Soma | $1e^{-1}$ - $4e^{-1}$ | 60 |
| Nav1.6 | Ais<br>Axon | $6e^{-3}$ - $8e^{-3}$<br>$6e^{-3}$ -$9e^{-3}$ | 60 |
| Kv3.4 | Soma<br>AIS<br>Axon | $5e^{-3}$ - $8e^{-3}$<br>$2e^{-2}$ - $4e^{-2}$<br>$1e^{-2}$ - $3e^{-2}$ | -84 |
| Kv4.3 | Soma<br>Dendprox | $4e^{-3}$ - $6e^{-3}$<br>$1e^{-3}$ - $4e^{-3}$ | -84 |
| Kv1.1 | Soma<br>AIS<br>Axon<br>Dendprox<br>Denddist | $6e^{-4}$ - $4e^{-3}$<br>$3e^{-4}$ - $5e^{-4}$<br>$2.5e^{-3}$ - $5e^{-3}$<br>$4e^{-3}$ - $1e^{-2}$<br>$1e^{-3}$ - $3e^{-3}$ | -84 |
| Kir2.3 | Soma | 1e-5 - 5e-5 | -84 |
| Kv7 | AIS | 6e-5 - 8e-5 | -84 |
| KCa1.1 | Soma<br>Dendprox<br>Denddist | $4e^{-3}$ - $9e^{-3}$<br>$1e^{-3}$ - $5e^{-3}$<br>$1e^{-3}$ - $4e^{-3}$ | -84 |
| KCa2.2 | Soma<br>Dendprox<br>Denddist | 4e-4 - 9e-4<br>3e-06 - 4.5e-06<br>1e-05 - 2e-05 | -84 |
| Cav2.1 | Soma<br>Dendprox<br>Denddist | $2e^{-4}$ - $3.5e^{-4}$<br>$4e^{-4}$ - $7e^{-4}$<br>2e-4 - $4e^{-4}$ | 137.5 |
| Cav3.2 | Soma<br>Dendprox | $9e^{-4}$ - $2e^{-3}$<br>$7e^{-4}$ - $1e^{-3}$ | 137.5 |
| Cav3.3 | Soma<br>Dendprox | $1.5e^{-05}$ - $2e^{-05}$<br>$1e^{-05}$ - $2e^{-05}$ | 137.5 |
| HCN1 | Soma<br>AIS<br>Axon | $2e^{-4}$ - $7e^{-4}$<br>$7e^{-4}$ - $1e^{-3}$<br>$6e^{-4}$ - $1e^{-3}$ | -34 |
| Leak | Soma<br>AIS<br>Axon | 3e-5 | -48 |

The table shows the main properties of ionic channels used in the SC models. For each ionic channel type, the columns specify the maximum ionic conductance ($G_i$-max), ionic channels reversal potential ($E_{rev}$). The corresponding gating equations were written either in Hodgkin-Huxley (HH) style or in Markovian style.

***Supplementary Table 2. Electrotonic compartments in stellate cell models***

| Compartment | 1 stellate cell | 2 stellate cell | 3 stellate cell | 4 stellate cell |
|---|---|---|---|---|
| Soma | 1 section; area 37.3 μm$^2$ | 1 section; area 31.1 μm$^2$ | 1 section; area 70.1 μm$^2$ | 1 section; area 31.1 μm$^2$ |
| Proximal dendrites | 14 sections | 7 sections | 5 sections | 18 sections |
| Distal dendrites | 90 sections | 35 sections | 85 sections | 44 sections |
| Total length dendrites | 1162.8 μm | 755.1 μm | 875.7 μm | 587.4 μm |
| AIS | 1 section; length 25.5 μm | 1 section; length 13.8 μm | 1 section; length 46.6 μm | 1 section; length 26.6 μm |
| Axon | 14 sections; Length 191.0 μm | 20 sections; Length 361.1 μm | 76 sections; Length 1218.0 μm | 31 sections; Length 551.4 μm |

The table shows the morphological analysis with NEURON software of the four morphologies used for the multi-compartment SC models. The table reports the sections of the multi-compartment SC model along with their number, their length and the soma area.

***Supplementary Table 3. Synaptic model parameters***

| | AMPA receptor | NMDA receptor | $GABA_A$ receptor |
|---|---|---|---|
| Gmax (mS/cm$^2$) | 2300 | 10000 | 1600 |
| P | 0.15 | 0.15 | 0.42 |
| τ facil (ms) | 10.8 | 5 | 0 |
| τ recov (ms) | 35.1 | 8 | 38.7 |

The table summarizes the parameters used for modeling the AMPA, NMDA and $GABA_A$ receptors (Nieus et al., 2006; Nieus et al., 2014; Bidoret et al., 2015).

***Supplementary Table 4, 5. Spike features***

Spontaneous firing

| | Clampfit EXP (n=9) | eFEL EXP (n=9) | eFEL MOD (n=4) |
|---|---|---|---|
| $AP_{Ampl}$ (mV) p=0.04 | 42.6 ± 3.6 | 37.3 ± 5.2 | 59.1 ± 6.3* |
| $AP_{AHP}$ (mV) p=0.24 | -50.4 ± 2.1 | -44.2 ± 4.0 | -42.7 ± 0.8 |
| $AP_{Thr}$ (mV) p=0.29 | -34.2 ± 1.1 | -31.8 ± 1.3 | -31.8 ± 0.6 |
| $AP_{HW}$ (ms) p=0.26 | 0.76 ± 0.05 | 1.0 ± 0.2 | 1.1 ± 0.04 |
| AP Freq (Hz) p=0.84 | 24.2 ± 2.1 | 25.1 ± 2.1 | 22.8 ± 3.3 |
| $V_m$ (mV) p=0.95 | -41.7 ± 1.8 | -41.2 ± 1.2 | -40.9 ± 0.8 |

Injected current (16 pA)

| | Clampfit EXP (n=5) | eFEL EXP (n=5) | eFEL MOD (n=4) |
|---|---|---|---|
| $AP_{Ampl}$ (mV) p=0.01 | 33.7 ± 4.8 | 32.8 ± 5.0 | 57.7 ± 5.7** |
| $AP_{AHP}$ (mV) p=0.55 | -46.4 ± 5.2 | -46.6 ± 5.6 | -39.6 ± 0.6 |
| $AP_{Thr}$ (mV) p=0.97 | -31.3 ± 2.1 | -30.3 ± 5.1 | -30.3 ± 0.6 |
| $AP_{HW}$ (ms) p=0.4 | 0.96 ± 0.16 | 0.98 ± 0.14 | 1.2 ± 0.1 |
| AP Freq (Hz) p=0.06 | 80.4± 6.9 | 79.4 ± 7.6 | 54.4 ± 6.3 |

The tables show exemplar values of features, obtained from experimental traces (n = 9 used for the spontaneous firing recordings and n = 5 used for the current injection experimental protocols) and from simulations (n = 4) using eFEL and Clampfit software.

### *SUPPLEMENTARY FIGURES*

#### *Supplementary Figure 1. Ionic currents in stellate cell model sections*

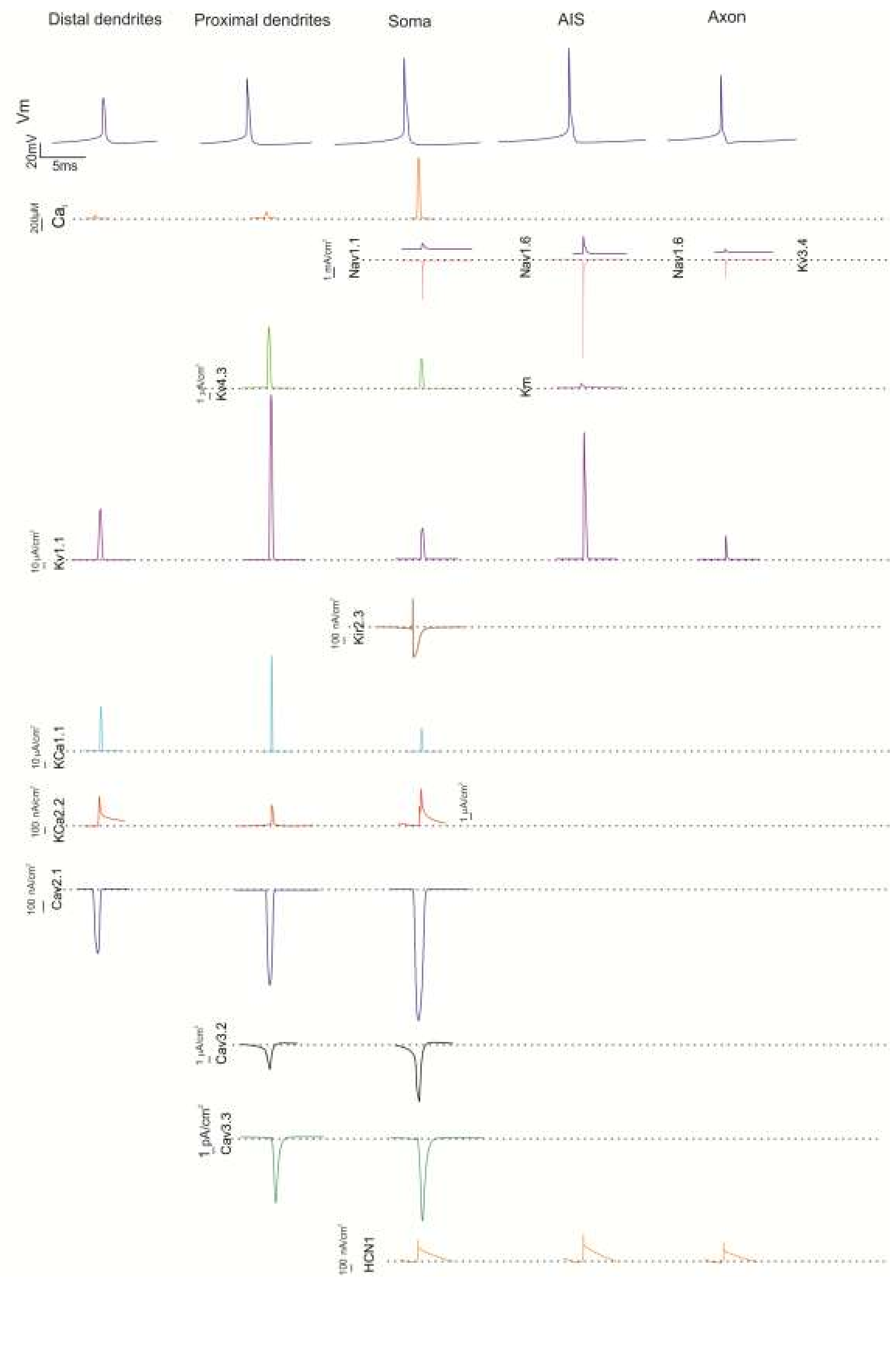

The traces show the ionic currents and calcium concentration changes generated by membrane channels in the SC model when a spike occurs during autorhythmic firing. Note the localization of channels in different sections and the different calibration scales.

***Supplementary Figure 2. AMPA-NMDA-GABA$_A$ receptors***

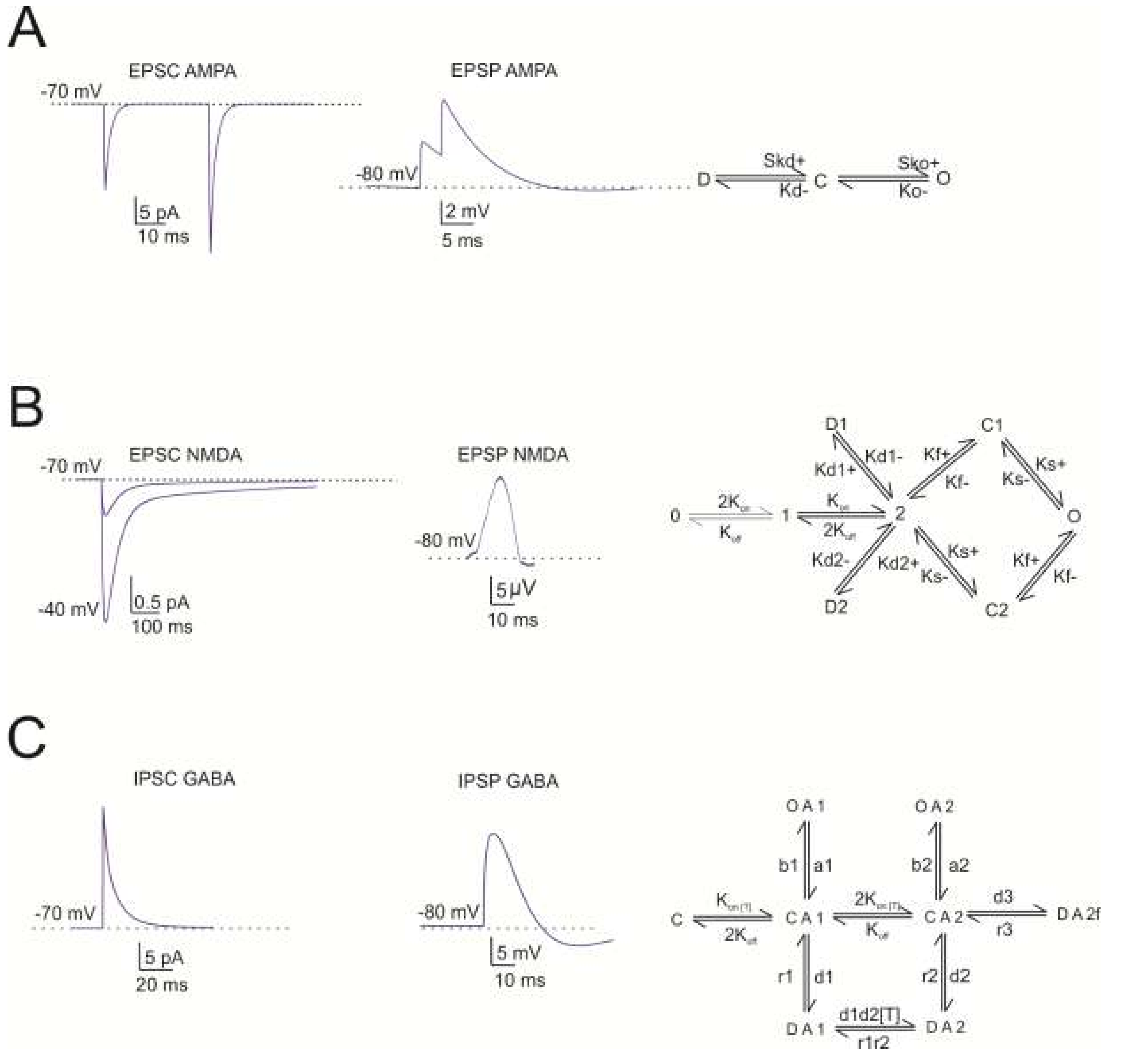


(A) The traces show simulated AMPA EPSC and EPSP. The AMPA receptor-channels kinetic scheme is shown on the left. (B) The traces show simulated NMDA EPSC and EPSP. The NMDA receptor-channels kinetic scheme is shown on the left. (C) The traces show simulated GABA-A IPSC and IPSP. The GABA-A receptor-channels kinetic scheme is shown on the left.

***Supplementary Figure 3. Ionic currents in the somatic compartment during responses to current injection***

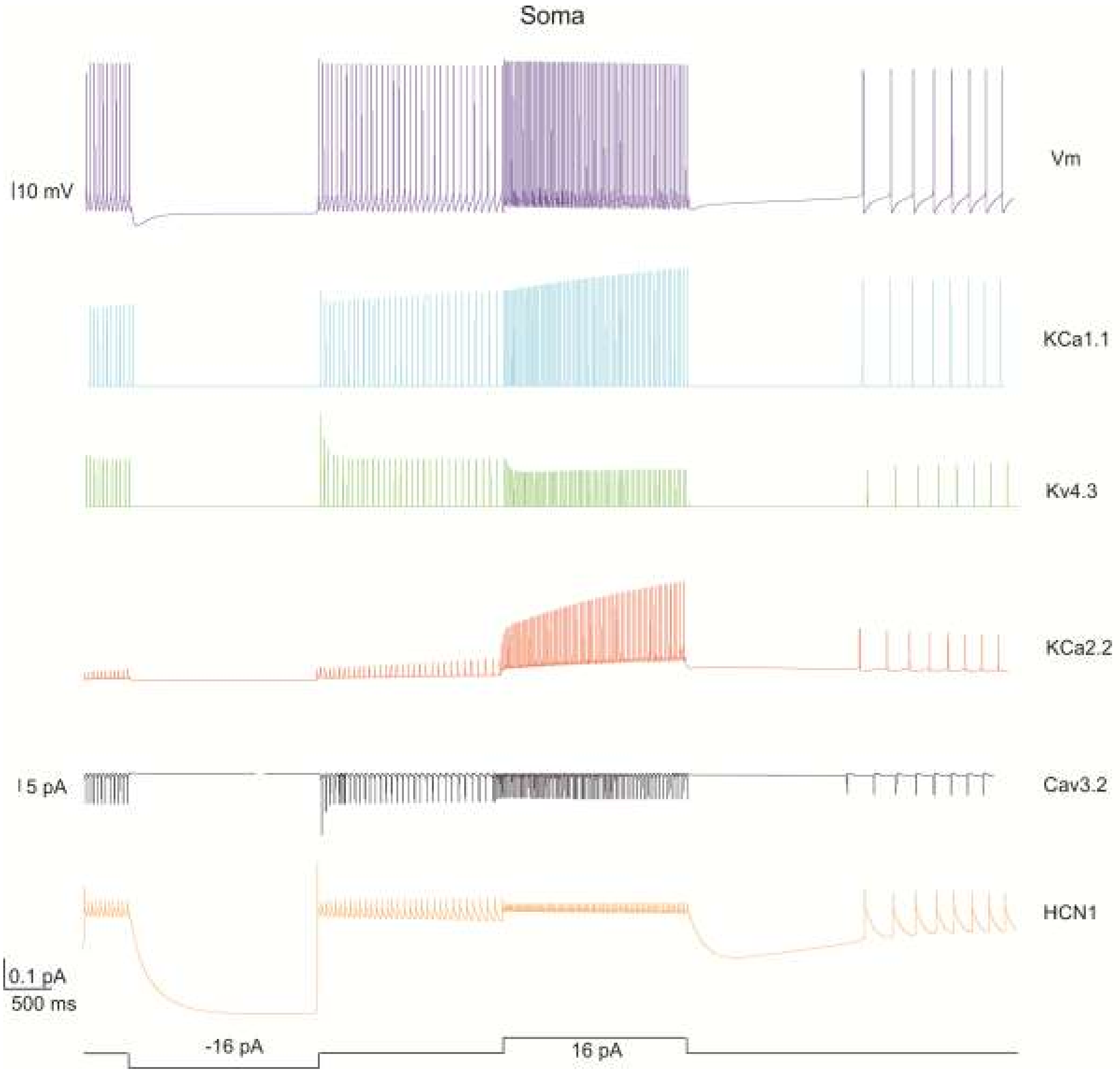


The traces show the model response during alternated phases of pacemaking, hyperpolarization and depolarization. The voltage trace ($V_m$) shows the membrane potential change. The current traces show (i) the dominant ionic currents in the soma involved in sagging inward rectification and rebound excitation in response to hyperpolarizing current injection (-16 pA), (ii) the dominant ionic currents in the soma in response to depolarizing current injection (16 pA).

*Supplementary Figure 4. Dendritic ionic currents during synaptic transmission*

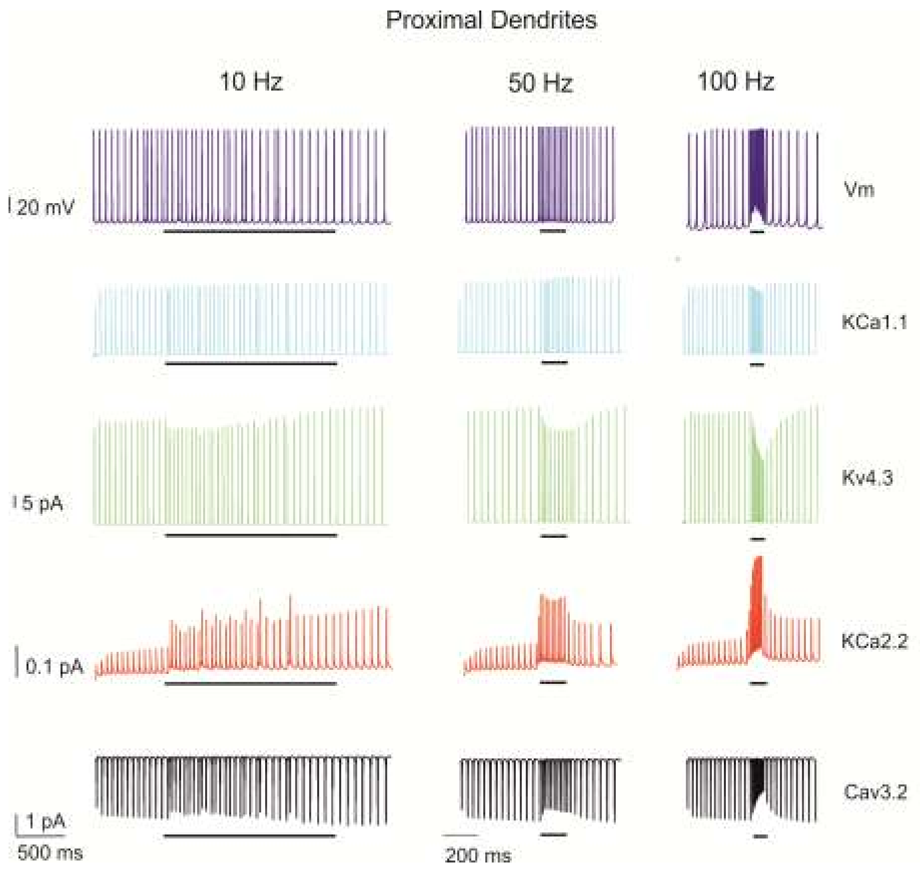


The traces show the SC model responses during a PF bursts of 10 impulses @ 10, 50 and 100 Hz. The voltage trace ($V_m$) shows the membrane potential change. The current traces show the dominant ionic currents in the proximal dendrites involved in the burst generation. Black bar indicates the stimulus duration.

*Supplementary Figure 5. Simulation of a long-duration burst*

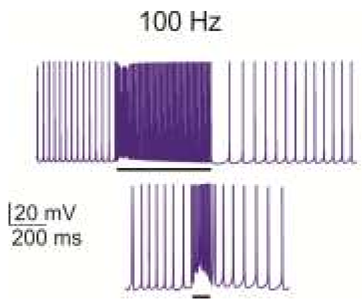


After a long duration burst (600 ms @ 100 Hz) a pause appears as with the injection of prolonged depolarizing currents. Black bar indicates the stimulus duration.

#### *SUPPLEMENTARY REFERENCES*


Akemann W, Knöpfel T (2006) Interaction of Kv3 potassium channels and resurgent sodium current influences the rate of spontaneous firing of Purkinje neurons. The Journal of neuroscience : the official journal of the Society for Neuroscience 26.

Alcami P, Marty A (2013) Estimating functional connectivity in an electrically coupled interneuron network. Proceedings of the National Academy of Sciences of the United States of America 110.

Anderson D, Engbers JD, Heath NC, Bartoletti TM, Mehaffey WH, Zamponi GW, Turner RW (2013) The Cav3-Kv4 complex acts as a calcium sensor to maintain inhibitory charge transfer during extracellular calcium fluctuations. J Neurosci 33:7811-7824.

Angelo K, London M, Christensen S, Häusser M (2007) Local and global effects of I(h) distribution in dendrites of mammalian neurons. The Journal of neuroscience : the official journal of the Society for Neuroscience 27.

Anwar H, Hong S, De Schutter E (2012) Controlling Ca2+-activated K+ channels with models of Ca2+ buffering in Purkinje cells. Cerebellum (London, England) 11.

Bastianelli E (2003) Distribution of calcium-binding proteins in the cerebellum. Cerebellum 2:242-262.

Bidoret C, Bouvier G, Ayon A, Szapiro G, Casado M (2015) Properties and molecular identity of NMDA receptors at synaptic and non-synaptic inputs in cerebellar molecular layer interneurons. Frontiers in synaptic neuroscience 7.

Brooke R, Atkinson L, Edwards I, Parson S, Deuchars J (2006) Immunohistochemical localisation of the voltage gated potassium ion channel subunit Kv3.3 in the rat medulla oblongata and thoracic spinal cord. Brain research 1070.

Brown D, Passmore G (2009) Neural KCNQ (Kv7) channels. British journal of pharmacology 156.

Cain S, Snutch T (2010) Contributions of T-type calcium channel isoforms to neuronal firing. Channels (Austin, Tex) 4.

Collin T, Cha tM, Lucas M, Moreno H, Racay P, Schwaller B, Marty A, Llano I (2005) Developmental changes in parvalbumin regulate presynaptic Ca2+ signaling. The Journal of neuroscience : the official journal of the Society for Neuroscience 25.

D'Angelo E, Nieus T, Maffei A, Armano S, Rossi P, Taglietti V, Fontana A, Naldi G (2001) Theta-frequency bursting and resonance in cerebellar granule cells: experimental evidence and modeling of a slow k+-dependent mechanism. J Neurosci 21:759-770.

Huguenard J, McCormick D (1992) Simulation of the currents involved in rhythmic oscillations in thalamic relay neurons. Journal of neurophysiology 68.

Indriati DW, Kamasawa N, Matsui K, Meredith AL, Watanabe M, Shigemoto R (2013) Quantitative Localization of Cav2.1 (P/Q-Type) Voltage-Dependent Calcium Channels in Purkinje Cells: Somatodendritic Gradient and Distinct Somatic Coclustering with Calcium-Activated Potassium Channels. In: J Neurosci, pp 3668-3678.

Kaufmann W, Kasugai Y, Ferraguti F, Storm J (2010) Two distinct pools of large-conductance calcium-activated potassium channels in the somatic plasma membrane of central principal neurons. In: Neuroscience, pp 974-986.

Khaliq Z, Gouwens N, Raman I (2003) The contribution of resurgent sodium current to high-frequency firing in Purkinje neurons: an experimental and modeling study. The Journal of neuroscience : the official journal of the Society for Neuroscience 23.

Kulik A, Nakadate K, Hagiwara A, Fukazawa Y, Luján R, Saito H, Suzuki N, Futatsugi A, Mikoshiba K, Frotscher M, Shigemoto R (2004) Immunocytochemical localization of the

alpha 1A subunit of the P/Q-type calcium channel in the rat cerebellum. The European journal of neuroscience 19.
Lorincz A, Nusser Z (2008) Cell-type-dependent molecular composition of the axon initial segment. The Journal of neuroscience : the official journal of the Society for Neuroscience 28.
Luján R, Albasanz J, Shigemoto R, Juiz J (2005) Preferential localization of the hyperpolarization-activated cyclic nucleotide-gated cation channel subunit HCN1 in basket cell terminals of the rat cerebellum. The European journal of neuroscience 21.
Magistretti J, Castelli L, Forti L, D'Angelo E (2006) Kinetic and functional analysis of transient, persistent and resurgent sodium currents in rat cerebellar granule cells in situ: an electrophysiological and modelling study. J Physiol 573:83-106.
Masoli S, Solinas S, D'Angelo E (2015) Action potential processing in a detailed Purkinje cell model reveals a critical role for axonal compartmentalization. Front Cell Neurosci 9:47.
Miceli F, Vargas E, Bezanilla F, Taglialatela M (2012) Gating currents from Kv7 channels carrying neuronal hyperexcitability mutations in the voltage-sensing domain. Biophysical journal 102.
Molineux ML, Fernandez FR, Mehaffey WH, Turner RW (2005) A-type and T-type currents interact to produce a novel spike latency-voltage relationship in cerebellar stellate cells. J Neurosci 25:10863-10873.
Molineux ML, McRory JE, McKay BE, Hamid J, Mehaffey WH, Rehak R, Snutch TP, Zamponi GW, Turner RW (2006) Specific T-type calcium channel isoforms are associated with distinct burst phenotypes in deep cerebellar nuclear neurons. Proc Natl Acad Sci U S A 103:5555-5560.
Nieus T, Sola E, Mapelli J, Saftenku E, Rossi P, D'Angelo E (2006) LTP regulates burst initiation and frequency at mossy fiber-granule cell synapses of rat cerebellum: experimental observations and theoretical predictions. J Neurophysiol 95:686-699.
Nieus TR, Mapelli L, D'Angelo E (2014) Regulation of output spike patterns by phasic inhibition in cerebellar granule cells. Front Cell Neurosci 8:246.
Pan Z, Kao T, Horvath Z, Lemos J, Sul J, Cranstoun S, Bennett V, Scherer S, Cooper E (2006) A common ankyrin-G-based mechanism retains KCNQ and NaV channels at electrically active domains of the axon. The Journal of neuroscience : the official journal of the Society for Neuroscience 26.
Perney T, Marshall J, Martin K, Hockfield S, Kaczmarek L (1992) Expression of the mRNAs for the Kv3.1 potassium channel gene in the adult and developing rat brain. Journal of neurophysiology 68.
Prüss H, Derst C, Lommel R, Veh R (2005) Differential distribution of individual subunits of strongly inwardly rectifying potassium channels (Kir2 family) in rat brain. Brain research Molecular brain research 139.
Raphemot R, Lonergan DF, Nguyen TT, Utley T, Lewis LM, Kadakia R, Weaver CD, Gogliotti R, Hopkins C, Lindsley CW, Denton JS (2011) Discovery, Characterization, and Structure–Activity Relationships of an Inhibitor of Inward Rectifier Potassium (Kir) Channels with Preference for Kir2.3, Kir3.X, and Kir7.1. Front Pharmacol 2.
Rehak R, Bartoletti TM, Engbers JDT, Berecki G, Turner RW, Zamponi GW (2013) Low Voltage Activation of KCa1.1 Current by Cav3-KCa1.1 Complexes. In: PLoS One.
Rowan M, Tranquil E, Christie J (2014) Distinct Kv channel subtypes contribute to differences in spike signaling properties in the axon initial segment and presynaptic boutons of cerebellar interneurons. The Journal of neuroscience : the official journal of the Society for Neuroscience 34.
Schaller K, Caldwell J (2003) Expression and distribution of voltage-gated sodium channels in the cerebellum. Cerebellum (London, England) 2.

Solinas S, Forti L, Cesana E, Mapelli J, De Schutter E, D'Angelo E (2007a) Computational reconstruction of pacemaking and intrinsic electroresponsiveness in cerebellar Golgi cells. Front Cell Neurosci 1:2.
Solinas S, Forti L, Cesana E, Mapelli J, De Schutter E, D'Angelo E (2007b) Fast-reset of pacemaking and theta-frequency resonance patterns in cerebellar golgi cells: simulations of their impact in vivo. Front Cell Neurosci 1:4.
Southan A, Robertson B (2000) Electrophysiological characterization of voltage-gated K(+) currents in cerebellar basket and purkinje cells: Kv1 and Kv3 channel subfamilies are present in basket cell nerve terminals. The Journal of neuroscience : the official journal of the Society for Neuroscience 20.
Stonehouse A, Pringle J, Norman R, Stanfield P, Conley E, Brammar W (1999) Characterisation of Kir2.0 proteins in the rat cerebellum and hippocampus by polyclonal antibodies. Histochemistry and cell biology 112.
Swensen AM, Bean BP (2003) Ionic Mechanisms of Burst Firing in Dissociated Purkinje Neurons. J Neurosci 23:9650-9663.
Turner R, Zamponi G (2014) T-type channels buddy up. Pflugers Archiv : European journal of physiology 466.
Williams M, Fuchs J, Green J, Morielli A (2012) Cellular mechanisms and behavioral consequences of Kv1.2 regulation in the rat cerebellum. The Journal of neuroscience : the official journal of the Society for Neuroscience 32.
Womack MD, Hoang C, Khodakhah K (2009) Large conductance calcium-activated potassium channels affect both spontaneous firing and intracellular calcium concentration in cerebellar Purkinje neurons. Neuroscience 162:989-1000.
Xu J, Clancy CE (2008) Ionic Mechanisms of Endogenous Bursting in CA3 Hippocampal Pyramidal Neurons: A Model Study. In: PLoS One.
